# A Blueprint for Broadly Effective Bacteriophage Therapy Against Bacterial Infections

**DOI:** 10.1101/2024.04.20.590411

**Authors:** Minyoung Kevin Kim, Qingquan Chen, Arne Echterhof, Robert C. McBride, Nina Pennetzdorfer, Niaz Banaei, Elizabeth B. Burgener, Carlos E. Milla, Paul L. Bollyky

## Abstract

Bacteriophage therapy is a tantalizing therapeutic option for anti-microbial resistant bacterial infections but is currently limited to personalized therapy due to the narrow host range of individual phages. Theoretically, cocktails incorporating numerous phages targeting all possible bacterial receptor specificities could confer broad host range. Practically, however, extensive bacterial diversity and the complexity of phage-phage interactions precludes this approach. Here, using screening protocols for identifying “complementarity groups” of phages using non-redundant receptors, we generate effective, broad-range phage cocktails that prevent emergence of bacterial resistance. Further, phage complementarity groups have characteristic interactions with particular antibiotic classes, making it possible to predict phage-antibiotic as well as phage-phage interactions. Using this strategy, we generate three phage-antibiotic cocktails, each effective against >96% of 153 *Pseudomonas aeruginosa* clinical isolates, including when used in biofilm cultures and wound infections *in vivo*. We similarly develop effective *Staphylococcus aureus* phage-antibiotic cocktails and demonstrate the utility of combined cocktails against polymicrobial (mixed *P. aeruginosa/S. aureus*) cultures, highlighting the broad applicability of this approach. These studies establish a blueprint for effective, broad-spectrum phage therapy cocktails and enable off-the-shelf phage-based therapeutics for antimicrobial-resistant bacterial infections.

## Introduction

The emergence of antimicrobial-resistant (AMR) pathogens urgently requires the development of novel therapeutic approaches to treat these bacterial infections^1^. There is great interest in using bacteriophages (phages) – viruses that kill bacteria – as alternatives or supplements to conventional antibiotics^2^. Indeed, phage therapy is already saving individual lives in the setting of compassionate use cases^3,4^. However, successful clinical trials and widely effective phages have been elusive^5–7^, limiting the therapeutic and commercial potential of this approach.

One major obstacle to effective phage therapy is the narrow host range of many individual phages. Although the level of specificity can vary, phages typically infect one particular host bacterial species and a limited set of strains (i.e., strains being variants within a single species). Bacterial receptors are implicated as major determinants of phage host range^8^, but the receptors for most phages remain unknown. Identifying phages with adequate coverage is particularly challenging in the setting of polymicrobial (made up of multiple microbial species) and polyclonal (made up of multiple bacterial strains) infections. For example, chronic wound infections in diabetic ulcers and chronic lung infections in cystic fibrosis often involve polyclonal infections with many distinct, AMR strains of *Pseudomonas aeruginosa*^9^ and also polymicrobial infections including both *P. aeruginosa* and *Staphylococcus aureus* simultaneously^10^. Identifying phages that can target all of these complex niches is an unsolved problem.

Another barrier is the development of phage resistance by host bacteria^11^. Bacteria employ diverse defense mechanisms to evade predation by phages^12,13^. Phage defense mechanisms are frequently triggered in response to phage predation, resulting in bacterial growth after initial inhibition^14^. Given these issues, laborious, time-consuming screening protocols are needed to identify effective phages for each bacterial isolate^15^.

“Cocktails” that combine multiple phages are an appealing solution to the problems of limited host range and the emergence of resistance^16,17^. However, phage-phage interactions can either be synergistic^18^ or antagonistic^19^ and are difficult to predict. Moreover, the optimum number of phages, and whether they ought to be administered serially or in concert are unclear. Further complicating the development of cocktail approaches, we lack the technical protocols, analytical frameworks, and appropriate pre-clinical models to effectively interrogate phage-phage and phage-antibiotic combinations.

Without clear guiding principles and effective protocols to inform cocktail design, phage cocktails are typically selected empirically^20,21^. Combining phages with antibiotics is similarly challenging, as phage-antibiotic interactions are also difficult to predict. The pairing of antibiotics with phages is likewise typically empiric^22–24^.

Genomic or strain-based approaches to improve phage cocktail design have been proposed^16,25^, but have proven difficult to implement clinically. Bacterial receptor diversity is an important determinant of phage host range^16,26^ and theoretically cocktails targeting all possible bacterial receptor specificities would have broad coverage. Similarly, matching phages to all of the individual bacteria in a population has shown promise against limited numbers of strains^27–29^, but these approaches are difficult to scale up to encompass the breadth of bacterial diversity. Unfortunately, current phage cocktails designs have shown limited success in clinical trials^5,24^ and the field has struggled to move past personalized phage therapy.

We hypothesized that identifying “complementarity” groups of phages using non-redundant receptors could be the key to generating effective phage-antibiotic cocktails for broad use. To test this idea, we developed an integrated series of assays and analytical approaches to facilitate cocktail selection. These efforts focused on two major pathogens commonly associated with devastating AMR infections^30,31^: *P. aeruginosa*, a Gram-negative bacterium, and *S. aureus*, a Gram-positive bacterium.

## Results

### Repeated exposure to phages promotes selection for heritable, high-level resistance

To study phage host range, we used a spectrophotometer to provide a robust, quantitative measurement of bacterial growth of *P. aeruginosa* strain PA14 by recording the optical density at a wavelength of 600 (OD600) over time^32^. Phage DMS3*vir* (a phage strain modified to be virulent^33^) was then introduced at a multiplicity of infection (MOI) of 100. We then assessed its impact on PA14 growth over 30 hours (**Supplementary** Fig. 1A). This time window was chosen because PA14 reaches a growth peak and is sensitive to phage predation within this time.

To quantify phage susceptibility from these growth measurements^34^, we calculated what we termed as the “Suppression Index”– the percentage of growth inhibition caused by phage within the first 30 hours of exposure (**Fig. 1A-i**). Initial bacterial growth is inhibited due to the predation by DMS3*vir*, but after 15 hours PA14 shows “growth”, indicating the fraction of phage-exposed bacteria develop resistance (**Fig. 1A-ii**). We obtained comparable results regardless of the MOI used, despite the sub-maximal initial inhibition effects at lower MOIs (**Fig. 1A-iii**). To maximize susceptibility to phages, we used an MOI of 100 throughout this work, unless indicated otherwise. Analogous results were obtained across a panel of eight different phages (**Fig. 1A-iv**).

**Figure 1.**
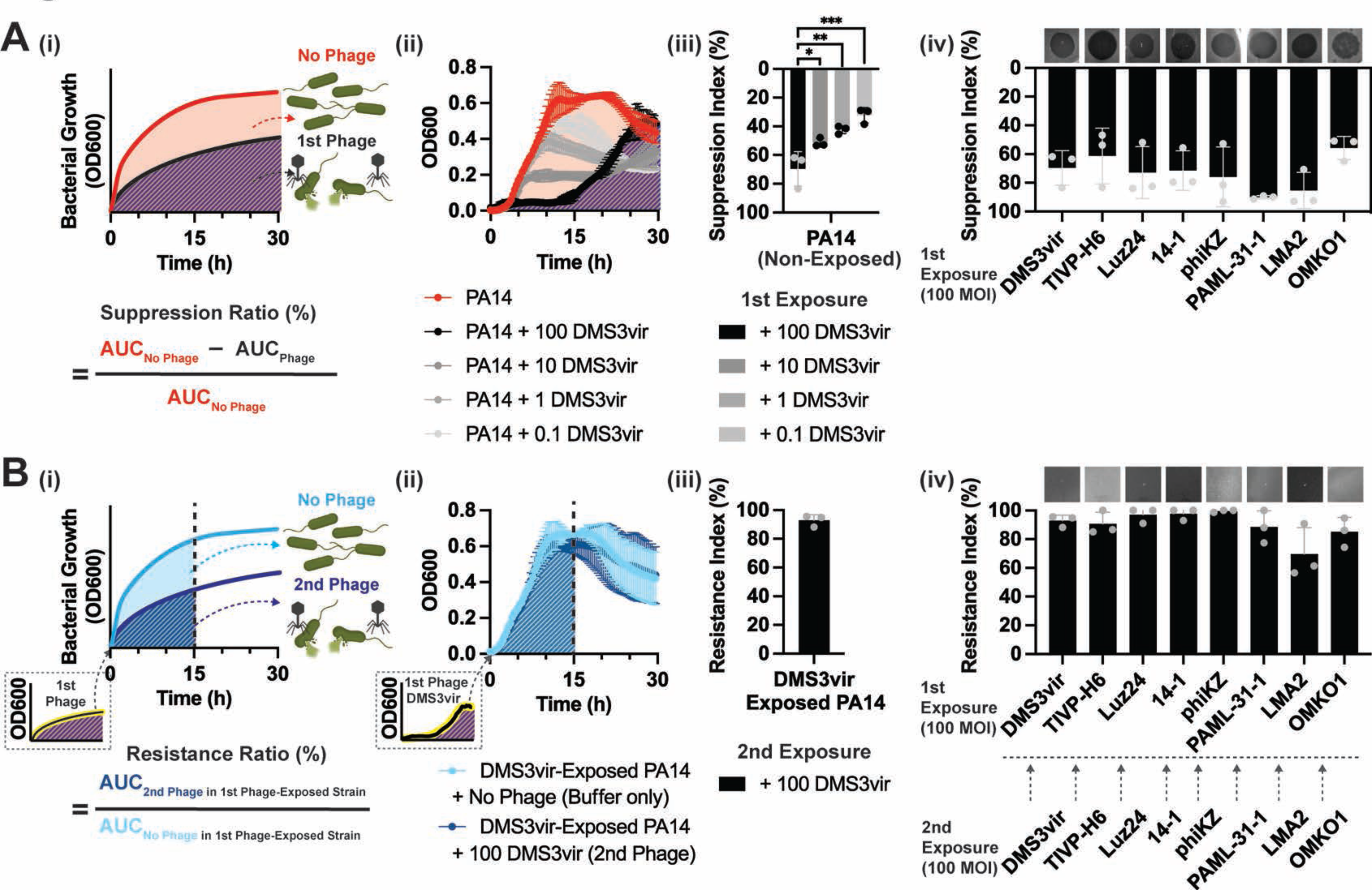
| Repeated exposure of bacteria to phages selects for heritable, constitutive resistance. (**A-i**), To quantify phage-mediated suppression of bacterial growth, we measured growth curves over 30 hours (h) and calculated the **“**Suppression Index”. This was defined as the area under the curve (AUC) for the non-phage-treated condition minus the phage-treated condition divided by the AUC for the non-treated condition. **(A-ii, A-iii),** Growth curves **(A-ii**) and Suppression Index **(A-iii)** for *P. aeruginosa* strain PA14 under different multiplicity of infection (MOI) of DMS3vir phage. **(A-iv),** Suppression Index for PA14 exposed to eight different phages, each at 100 MOI. Representative plaque assays from triplicate experiments are shown above each column. **(B-i),** To quantify resistance upon repeat phage exposure, we collected bacteria following an initial phage exposure (1st exposure) and then re-cultured these bacteria in the absence or presence of the identical phage (2nd exposure). Upon this second exposure, we again generated growth curves over 15 h and determined the “Resistance Index”, defined as the AUC of the phage-rechallenged condition divided by the AUC of the non-rechallenged condition. **(B-ii, B-iii)**, Growth curves (**B-ii**) for PA14 after initial exposure to 100 MOI of DMS3vir (1st exposure) followed by subsequent exposure to either 100 MOI DMS3vir (2nd exposure, dark blue) or Buffer control (light blue) and the resulting Resistance Index (**B-iii**). (**B-iv**), Resistance Index for PA14 challenged initially with one of eight different phages (1st exposure) and subsequently re-challenged with the same phage (2nd exposure). Representative plaque assays are shown above each column. All error bars denote standard deviations from at least triplicate experiments. *P <0.05, **P<0.005 and ***P <0.0005 for multiple unpaired t-tests.

We then re-exposed these same bacterial cultures a second time to the same phages (**Fig. 1B-i**). We observed that PA14 variants occasionally emerged with a complete loss of susceptibility to the phage (**Fig. 1B-ii**). Following the initial phage exposure, this loss of susceptibility was constitutive, akin to a loss of host range^35^.

To quantify this phage resistance upon repeated exposure, we calculated what we call the “Resistance Index” – the percentage of growth observed over 15 hours upon phage re-challenge in a previously phage-exposed strain (**Fig. 1B-i**). For example, we observed more than 90% bacterial growth (or 90.1% Resistance Index), when re-exposed to 100 MOI of DMS3*vir* in the previously DMS3*vir*-exposed strain (**Fig. 1B-ii, 1B-iii)**, indicating that the previously DMS3*vir*-exposed strain had developed near-complete resistance to DMS3*vir*. Similar results were obtained across a panel of eight different phages (**Fig. 1B-iv**).

### Constitutive phage resistance confers cross-resistance against “complementarity groups” (CGs) of genetically diverse phage species

Next, we asked how constitutive phage resistance induced upon repeated exposure to one phage affects susceptibility to other phages. To this end, each phage-exposed PA14 strain was “rechallenged” by all eight phages, and the individual Resistance Indexes were assessed. We observed that bacteria that lost susceptibility to each of the three phages DMS3*vir*, TIVP-H6, and Luz24 lost susceptibility to all three phages in this group. Similarly, bacteria that lost susceptibility to either 14-1 or PhiKZ lost susceptibility to both (**Fig. 2A**). This cross-resistance within a group of phages indicated a shift in host range, as similarly shown previously^36^.

**Figure 2.**
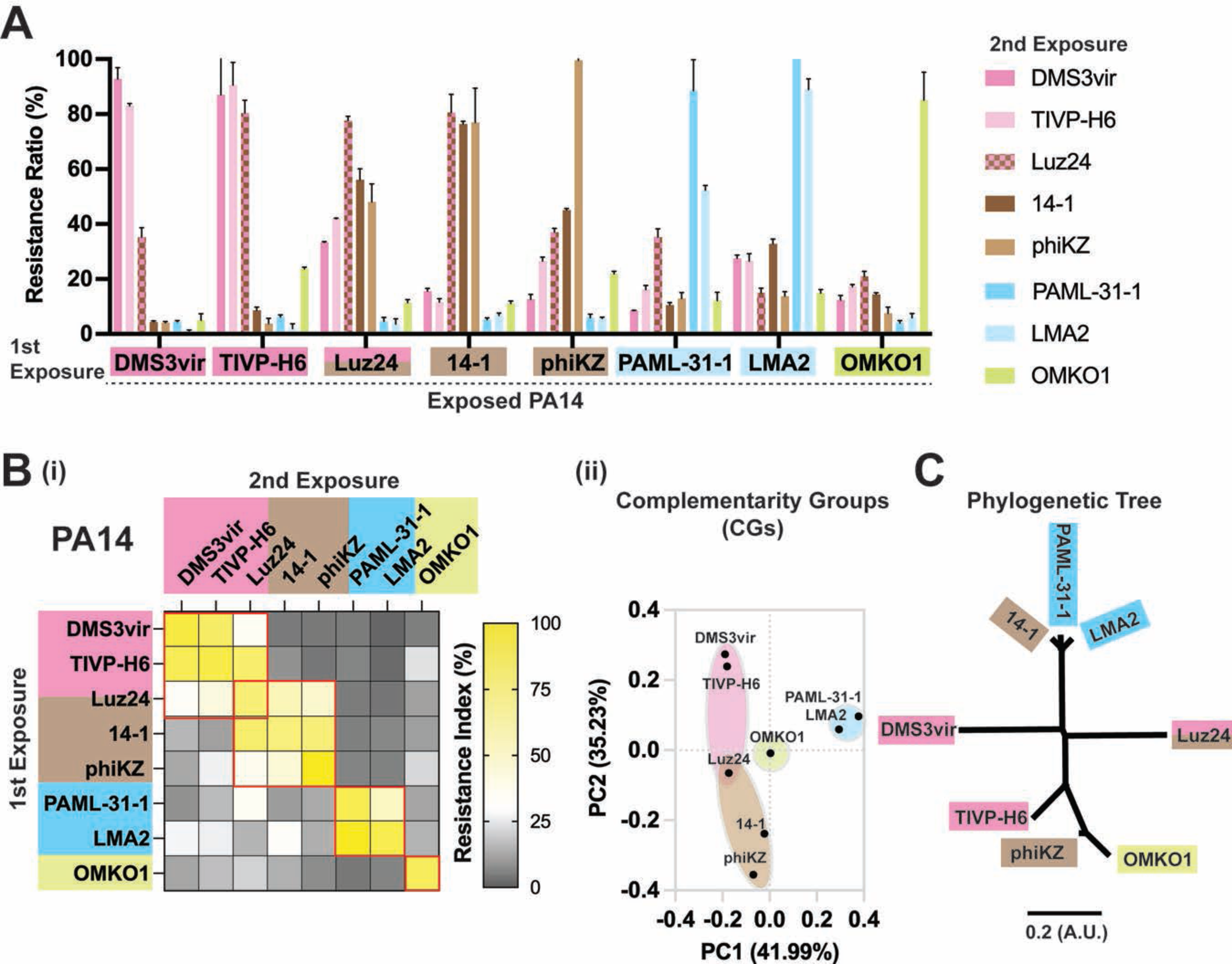
| Constitutive phage resistance confers cross-resistance against “complementarity groups” (CG) of genetically diverse phages. **A**, Resistance Index of PA14 initially exposed to one of a set of eight phages and then subsequently re-exposed to each of the same set of eight phages, all with MOI = 100. Error bars denote standard deviations from triplicate experiments. (**B-i**), Phage exposure matrix representing the average value of the Resistance Index data presented in Figure 2A. These data highlight the presence of multiple groups of phages wherein constitutive resistance to one confers PA14 cross-resistance against other phages within that group. Here, CGs are defined as groups of phages that can have cross-constitutive resistance to one another within that group. (**B-ii**), PCA analysis of the chart description of (**B-i**). **C**, Phylogenic dendrogram for the phages in panels **A, B**. The scale (A.U.) is represented as relative evolutionary distance (substitution/site).

Using these cross-resistance patterns, we generated a phage exposure matrix (**Fig. 2B-i**). We observed that phages clustered within four major groups, which we termed “complementarity groups” or CGs. These phage CG patterns are further supported by principal component analysis (PCA) (**Fig. 2B-ii),** which shows the phages with the same CG clustered with one another. Notably, phage Luz24 belongs to two CGs, indicating that dual identity is possible. Analogous CG results were observed using *P. aeruginosa* strain PAO1 (**Supplementary** Fig. 2A-i**,ii**), suggesting that phage CG structures are generalizable across *P. aeruginosa* strains.

Next, we investigated whether the resistance matrix chart could be expanded to include previously uncharacterized phages. To this end, we isolated a pair of novel *P. aeruginosa* phages, which we called KOR_P1 and KOR_P2, from local sewage (**Supplementary** Fig. 2B-i**,ii**). The newly identified phages could likewise be linked to phage susceptibility groups based on cross-resistance patterns (**Supplementary** Fig. 2B**-iii**). Upon screening and analysis, we observed that both KOR_P1 and KOR_P2 grouped with the CG of DMS3*vir* (**Supplementary** Fig. 2B**-iv**), suggesting that the CG framework is expandable.

We also questioned whether CG patterns might be correlated with the evolutionary relationships among these phage species^8^. To test this, we generated a phylogenetic tree based on phage genome sequences. We observed that the phylogenetic grouping did not entirely align with the CG patterns (**Fig. 2C)**, indicating that the determinants of CG may not necessarily be evolutionarily fixed nor mediated by convergence evolution, presumably due to the mosaicism nature of phage genome^37^.

### Phages belonging to the same CG use the same bacterial receptors

We hypothesized that phages within a single CG share the same bacterial receptors, such that loss of susceptibility to one phage confers loss of susceptibility to all. In support of this model, phages are known to use common bacterial structures, including the Type IV pilus, the flagella, and oligosaccharide antigens/lipopolysaccharide (OSA/LPS)^38–41^, as receptors for entry to their hosts (**Fig. 3A-i)**.

**Figure 3.**
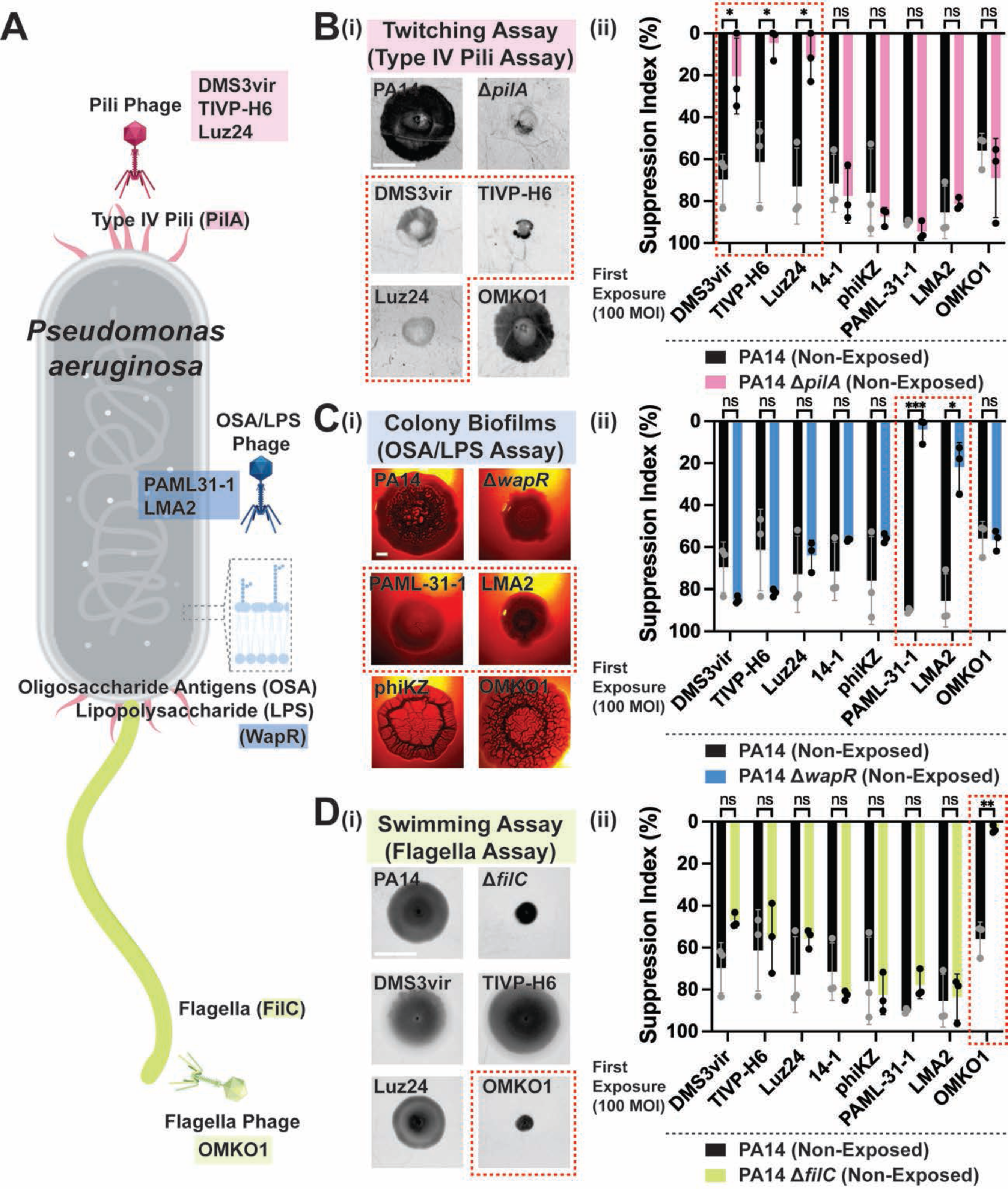
| Phages belonging to the same CG use the same bacterial receptors. **A**, Schematic model description of three major surface receptors involved in phage uptake by *P. aeruginosa*, including Type IV pili (pink), oligosaccharides antigens (OSA) and lipopolysaccharide (LPS, blue), and flagella (green). (**B-i)**, Representative images of twitching motility assay by PA14, PA14Δ*pilA*, and DMS3vir-, TIVP-H6-, Luz24-, and OMKO1-exposed PA14. **(B-ii),** Suppression Index measured for PA14 (black) and the PA14Δ*pilA* (pink) by eight different types of phages, each with MOI = 100. **(C-i),** Colony biofilm phenotypes of PA14, PA14Δ*wapR*, and the designated phage-exposed strains on Congo red agar medium after 100 h of growth. **(C-ii),** Suppression Index measured for the wild-type PA14 (black) and PA14Δ*wapR* (blue) by eight different types of phages at MOI = 100. **(D-i),** Representative images of swimming motility assay by PA14, PA14Δ*filC*, and DMS3vir-, TIVP-H6-, Luz24-, and OMKO1-exposed PA14. **(D-ii),** Suppression Index measured for PA14 (black) and PA14Δ*filC* (green) by eight different types of phages, each at MOI = 100. Error bars denote standard deviations from triplicate experiments. *P < 0.05, **P<0.005 and ***P < 0.0005 for multiple unpaired t-tests. Images are based on a minimum of triplicate independent replicates. Scale bars = 10 mm.

To investigate this idea, we asked whether resistance to a particular phage CG could result in a phenotype consistent with the loss of a functional receptor. We found that PA14 that lost susceptibility to either DMS3*vir*, TIVP-H6, or Luz24 phage CG no longer exhibited twitching motility, known to be mediated by the Type IV pilus, and resembled a Type IV pilus mutant strain (PA14Δ*pilA*)(**Fig. 3B-i; Supplementary** Fig. 3A-i). To further support this notion, we found that the phages in this CG require an intact Type IV pilus to infect PA14 (**Fig. 3B-ii; Supplementary** Fig. 3A**-ii)**. We obtained identical results – reduced infection by the DMS3*vir*, TIVP-H6, and Luz24 phages – with seven different mutant strains defective in various components of the type IV pilus, including PA14Δ*pilA*, PA14Δ*pilB*, PA14Δ*pilC*, PA14Δ*pilE*, PA14Δ*pilX*, PA14Δ*pilX*, and PA14Δ*pilY1*, suggesting that DMS3*vir*, TIVP-H6, and Luz24 requires an intact Type IV pilus to infect PA14 (**Supplementary** Fig. 3A**-iii)**. Of note, DMS3*vir*, TIVP-H6, and Luz24 phages are still able to partially infect the PA14Δ*pilTU* mutant that has an intact Type IV pilus structure but has defective retraction motor proteins^42^, indicating that these phages did not need a fully functional exterior Type IV pilus structure to infect their targets. To further support the role of the Type IV pilus in infection by phages in this CG, we compared the whole genome sequencing (WGS) data of those phage-resistant PA14 strains upon phage exposure with the non-exposed PA14. We found that they had mutations in Type IV pilus genes (**Supplementary Table 1**), indicating that the loss of susceptibility to infection by phages in this CG is mediated by mutations in the genes harboring for Type IV pilus.

We also observed that the PA14 strains that had lost susceptibility to either LMA2 or PAML-31-1 exhibited a loss of rugae in the center of a colony biofilm, a phenotype generally associated with oligosaccharide antigen (OSA)-containing LPS, and instead resembled an OSA/LPS mutant strain of PA14 (PA14Δ*wapR*)(**Fig. 3C-i; Supplementary** Fig. 3B-i). The phenotypic changes in the PA14 strains that had lost susceptibility to either LMA2 or PAML-31-1 were also supported by a shift of OSA/LPS size in those strains (**Supplementary** Fig. 3B**-ii)**. These phages require an intact OSA/LPS to infect PA14 (**Fig. 3C-ii)**. Consistent with this finding, we compared the WGS data from PA14 strains that lost susceptibility to either LMA2 or PAML-31-1 upon phage exposure with the non-exposed PA14 strain, which revealed that they acquired mutations of genes that govern the synthesis of OSA/LPS (**Supplementary Table 1**). Together, these data indicate that the loss of PA14 susceptibility to infection by phages in this CG is mediated by mutations in the genes governing OSA/LPS synthesis.

Similarly, we observed that the PA14 strain that lost susceptibility to the phage OMKO1 diminished its swimming motility and resembled a strain of PA14 bearing mutant flagella (PA14Δ*filC*) (**Fig. 3D-i; Supplementary** Fig. 3C-i). We also found that this phage requires intact flagella to infect PA14, as evidenced by the phage’s inability to infect mutant strains defective in different components of the flagella structure, FilC (PA14Δ*filC*) and FlgK (PA14Δ*flgK*)(**Fig. 3D-ii; Supplementary** Fig. 3C**-ii, iii).** Of note, OMKO1 phage is also unable to infect the PA14Δ*oprM* mutant that has a defective outer membrane protein for antibiotic efflux^43^ (**Supplementary** Fig. 3C**-iv),** indicating that both intact flagella and OprM are required for OMKO1 to infect PA14. Consistent with these findings, we compared the WGS data from the PA14 strain that had lost susceptibility to OMKO1 upon phage exposure with the non-exposed PA14 strain, which revealed that it had mutations of genes for flagella and/or OprM (**Supplementary Table 1**). Together, these data indicate that the loss of PA14 susceptibility to infection by the phage OMKO1 in this CG is mediated by mutations in flagella and/or OprM.

We also investigated what receptor-related phenotype changes are involved in PA14 with resistance to phages PhiKZ and 14-1. However, while possible receptors for these phages have been suggested^44–46^, a clear receptor-related phenotype was not identified in this present work.

We were intrigued to note that some phages may belong to multiple complementarity groups. For example, phage Luz24 clearly requires the Type IV pilus for infectivity of PA14 (**Fig. 3B-i,ii)**, but phenotypically it also shares cross-resistance patterns with phages PhiKZ and 14-1 (**Fig. 2A, 2B).** Consistent with this result, this Luz24 phage has been reported previously to utilize either a cell surface structure^44^ or Type IV pilus as its entry receptors.

Together, these data indicate that a loss of susceptibility to a CG of phages is associated with loss/changes in expression of the entry receptor, which confers cross-resistance against the entire group of phages that utilize that receptor(s). This idea is further supported by our finding that the two newly identified phages (KOR_P1 and KOR_P2), which we found belong to the same CG as DMS3*vir* (**Supplementary** Fig. 2B**-iii**), also rely on the Type IV pilus for phage entry, as seen in studies with the *PA14*Δ*pilA* strain (**Supplementary** Fig. 3D). These data demonstrate that information on CG identity can help predict the receptor(s) used by newly identified phages.

### Cocktails containing phages from multiple CG reliably eliminate multi-drug resistant clinical isolates of *P. aeruginosa,* including biofilm and polyclonal cultures

We next sought to determine how many phages are optimal and whether combinations of phages from the same or different CGs were equally effective. To this end, we compared one phage versus two phages from the same CG, two phages from different CGs, or three phages from different CGs. We observed that while treatment with one or two phages from the same CG could initially suppress bacterial growth, treatment with two or three phages from different CGs more completely eliminated regrowth within the 30-hour time frame examined here (**Fig. 4A-i**). Pooled Suppression Index data for PA14 treated with various phage cocktails, each using a different combination of phages (**Fig. 4A-ii; Supplementary Table 2**), showed that two or more phages from different CGs can suppress bacterial growth by over 90%.

**Figure 4.**
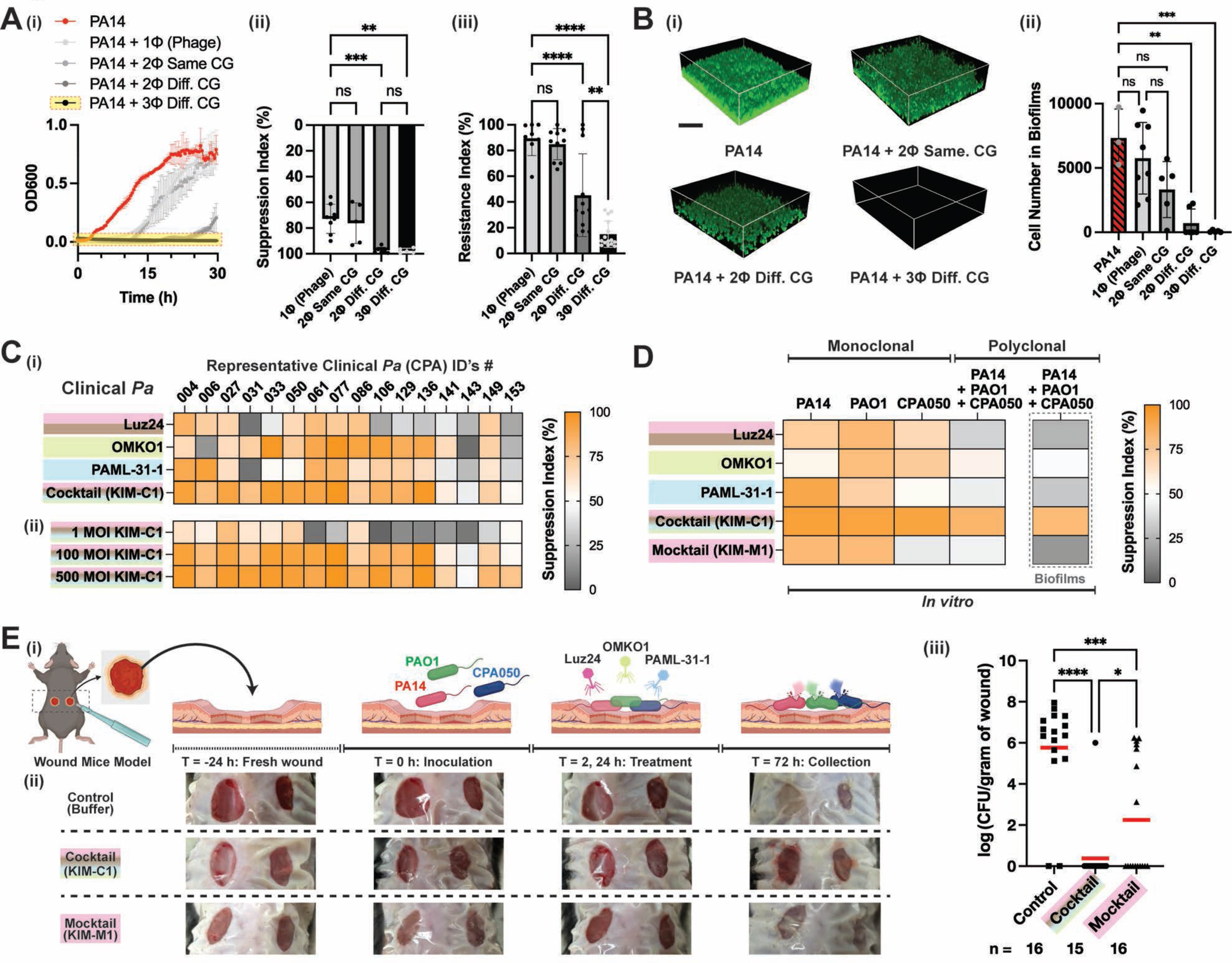
| Cocktails containing phages from three or more CG reliably eliminate multi-drug resistant clinical isolates of *P. aeruginosa,* including biofilm and polyclonal cultures *in vivo*. (**A-i**), Growth curves for PA14 treated with phage(s) from DMS3vir, the same CG (DMS3vir+TIVP-H6), two different CGs (DMS3vir+OMKO1), or three different CGs (Luz24+OMKO1+PAML-31-1). Of note, the treatment with three phages from different CGs completely prevented growth (highlighted in yellow). **(A-ii),** Pooled Suppression Index data for PA14 treated with different phage cocktails; each dot is represented as an average value of the Suppression Index from a different combination of phages with the number of phages and number of CGs as indicated. These cocktails are listed in Supplementary Table 2. **(A-iii)**, Pooled Resistance Index for PA14 treated initially with different phage cocktails and subsequently re-challenged by each component phage that comprises the cocktails, all at 100 MOI. Detailed information on the phages used is in Supplementary Table 2. **(B-i),** Three-dimensional renderings of PA14 biofilms after 72 hours in the absence or presence of phage therapy, as indicated. (**B-ii**), Number of PA14 cells in biofilms under the conditions in **B-i**. **(C-i),** Suppression Index for 16 clinical isolates of *Pa* by individual phages Luz24, OMKO, and PAML-31-1, each at 100 MOI, or a cocktail of these three phages (KIM-C1) with 100 combined MOI. **(C-ii)** Suppression Index for these clinical isolates of *P. aeruginosa* by cocktail KIM-C1 at 1, 100, or 500 combined MOIs. Data are shown as an average of triplicate independent results. **D,** Suppression Index for monoclonal and polyclonal *P. aeruginosa* cultures treated individually with KIM-C1 cocktail, its three constituent phages, or KIM-M1 mocktail (a mixture of 3 phages with same CG = DMS3vir, KOR_P1, and KOR_P2) grown in planktonic form or as biofilms. Data are shown as an average of triplicate independent results. **(E-i),** A schematic for wound mice model. **(E-ii),** Representative wound images of each condition (control, cocktail, and mocktail) over the course of experiments. **(E-iii),** CFU of the polyclonal cultures in wounds under the conditions in **E-ii**. The red line indicates the average value of each condition. “LOD” denotes the limit of detection. *P < 0.05, ***P<0.0005, and ****P < 0.00005 for multiple unpaired t-tests.

We then examined the development of constitutive resistance to phages upon repeated exposure to these four conditions. The pooled Resistance Index data for PA14 treated with various phage cocktails showed that three or more phages from different CGs most effectively suppressed resistance (**Fig. 4A-iii; Supplementary Table 2**).

Similarly, we found that three phages from different CGs were the most effective against biofilm cultures, as measured by confocal microscopy^47^ **(Fig. 4B-i, ii**). Together, these data demonstrated the superiority of combining three phages from different CGs in regard to suppressing regrowth without the development of resistance.

We next asked whether phages should be given in sequence or in concert. To this end, we compared sequential monotherapy versus consecutive dual therapy, as per the schematic in **Supplementary** Fig. 4A-i. We observed that phages delivered in combination suppressed bacteria growth over time with less resistance, as opposed to the phages delivered in series (**Supplementary** Fig. 4A**-ii).** Analogous results were obtained using other combinations of phages with different complementarity groups (**Supplementary** Fig. 4A**-iii, iv)**, suggesting that this is a general feature of phage CG cocktails. Together, these data indicate that phage cocktails given in concert are superior to those given in sequence, as suggested previously^19^.

We then examined the impact of phage cocktails incorporating multiple CGs on de-identified clinical *P. aeruginosa* isolates collected at Stanford University Medical Center. We observed that individual phages Luz24, OMKO, and PAML-31-1 were sporadically effective at suppressing the growth of 16 clinical isolates of *P. aeruginosa*. In contrast, a cocktail of these three phages (which we call “KIM-C1”) was more reliably effective **(Fig. 4C-i)**. Of note, we chose this particular three-phage combination as it covers all four different CGs. We also noticed that the effectiveness of KIM-C1 is also dose-dependent **(Fig. 4C-ii)**. These results indicate that the KIM-C1 phage cocktail can effectively target clinical isolates.

To explore the possibilities of this approach in microbially complex bacterial niches, we asked whether our strategy was also effective against mixed *P. aeruginosa* strains, such as might be found in a polyclonal infection^9^. To interrogate this, we grew three *P. aeruginosa* strains with comparable growth rates (PA14-mCherry, PAO1-GFP, and a clinical isolate of CPA050) together (**Supplementary** Fig. 4B**-i**) and then confirmed that by tracking labeling of each strain they do not interfere each other’s growth over 30 hours (**Supplementary** Fig. 4B**-ii**). We then treated this polyclonal culture with either Luz24, OMKO, and PAML-31-1 individually or a “cocktail” of these three phages administered together (KIM-C1). Once again, this cocktail covered four CGs. As a control, a “mocktail” of three phages (DMS3*vir*, KOR_P1, and KOR_P2) from a single CG was used and which we call “KIM-M1”. We observed that the individual *P. aeruginosa* strains are often susceptible to individual phages or the mocktail. However, the polyclonal culture was only eliminated by KIM-C1. This was also true for the same bacteria grown under biofilm conditions (**Fig. 4D**). These results indicate that polyclonal bacterial infections can be effectively targeted by a cocktail of phages from multiple CGs.

Next, we investigated whether our strategy is also applicable in an animal model of wound infection^48,49^. A schematic of this model is shown in **Fig. 4E-i**. In brief, we mixed equal portions of three *P. aeruginosa* strains (PA14-mCherry, PAO1-GFP, and CPA050) and infected mice with the mixed culture 24 hours after a fresh wound was made (**Fig. 4E-ii**). Then, each wound was treated twice (at 2 and 24 hours after bacterial infection) with either a cocktail (KIM-C1), a mocktail (KIM-M1), or a buffer as a control. At 72 hours after infection, we collected the wound and measured the remaining bacteria in each wound. We observed that the polyclonal culture was more consistently eradicated by the cocktail KIM-C1, as compared to the mocktail or control (**Fig. 4E-iii**). These results confirm that the phage cocktail, KIM-C1 can successfully treat polyclonal infections *in vivo*.

Together, our findings indicate that combinations of three or more phages from different CGs are effective at suppressing the growth of clinical *P. aeruginosa* isolates, as shown here in planktonic conditions, biofilms, polyclonal cultures, and in an *in vivo* animal model.

### Phage CGs have predictable interactions with particular classes of conventional antibiotics

Given that antibiotics are known to impact phage lytic activity in both positive (synergistic) and negative (antagonistic) ways^22,24,43,46,50^, we next sought to determine whether phages from particular complementarity groups similarly have characteristic relationships with different conventional antibiotics. To investigate this, we used low levels of antibiotic (sub-MIC levels) to best study phage/antibiotic interactions. Growth curves from PA14 under the different sub-MICs of 12 antibiotics used in these experiments are depicted in **Supplementary** Fig. 5A. Similarly, we used a low MOI (MOI = 10) of phages in these experiments. Then, we measured the bacterial growth over 30 hours in the presence of either a particular antibiotic, each phage, or both together (**Supplementary** Fig. 5B). The synergy score index was calculated as the antibiotic-phage interactions in each pair using the Highest Single Agent (HSA) Model, as detailed in the Methods section.

We observed that phages belonging to the same complementary group generally share common patterns of interactions with conventional antibiotics (**Fig. 5A)**. Some antibiotic classes were synergistic with most phages, such as the β-lactam drugs (e.g., ceftazidime (CTZ) and meropenem (MER)), while other antibiotic classes had idiosyncratic patterns of activity with phage GCs (e.g. ciprofloxacin (CIP), and doxycycline (DOX)). These data suggest that for some antibiotics, characteristic patterns of antibiotic responses exist for different phage complementarity groups.

**Figure 5.**
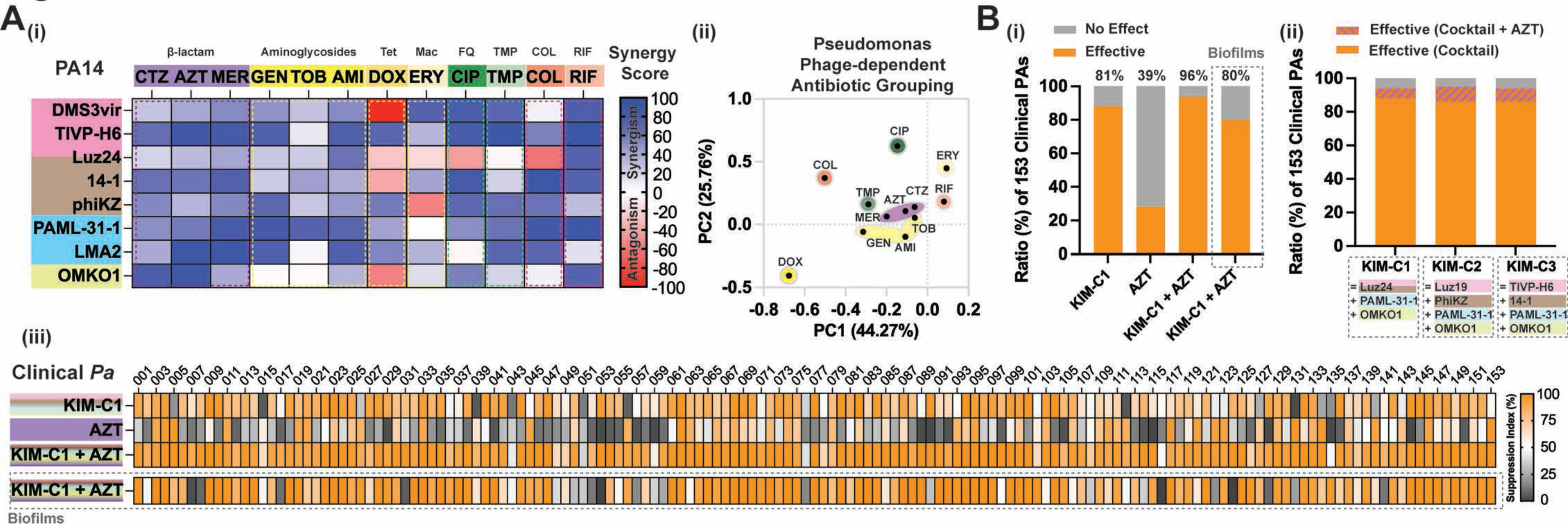
| Phage CGs have predictable, class-dependent interactions with conventional antibiotics. (**A-i**), Exposure matrix for PA14 exposed to combinations of phages and antibiotics simultaneously. Blue areas represent synergy while red areas represent antagonism. The antibiotic concentrations were below their MIC values (sub-MIC) as detailed in the Supplementary Information. CTZ = ceftazadime, AZT = aztreonam, MER = meropenem, GEN = gentamicin, TOB = tobramycin, AMI = amikacin, DOX = doxycycline, ERY = erythromycin, CIP = ciprofloxacin, TMP = trimethoprim, COL = colistin, and RIF = rifampin. Data are shown in the matrix as an average of triplicate independent results. **(A-ii),** PCA data for responses to phages and antibiotics. **(B-i),** Effectiveness Ratio (%) of 153 clinical *P. aeruginosa* isolates treated with the KIM-C1 cocktail (Luz24, OMKO, and PAML-31-1), AZT treatment alone, or the KIM-C1 cocktail plus aztreonam (KIM-C1+AZT) grown in planktonic form or as a biofilm. Effectiveness was defined as >60% growth suppression over 30 hours. **(B-ii),** Effectiveness Ratio (%) of 153 clinical *P. aeruginosa* isolates treated with one of three different phage cocktails (KIM-C1, –C2, and –C3), each with three or four phages from all four different CGs. (**B-iii**), Suppression Index for 153 clinical *P. aeruginosa* isolates treated with the KIM-C1 cocktail, AZT alone, or the KIM-C1 cocktail plus aztreonam (KIM-C1+AZT) grown in planktonic form or as biofilms. Data are shown in the matrix as an average of triplicate independent results.

To investigate the impact of combining antibiotics with our cocktail (shown in **Fig. 4**), we expanded our analysis to include 153 *P. aeruginosa* clinical isolates. We treated these with the cocktail KIM-C1, aztreonam (AZT), or a phage/antibiotic cocktail (KIM-C1+AZT) and then measured the Suppression Index for each. We observed comparable rates of effectiveness regardless of whether >50, >60, or >70% suppression was used as the cut-off for effectiveness (**Supplementary** Fig. 5C). For subsequent analyses, >60% suppression was used as the cut-off for effective therapy in these studies.

We observed that among 153 clinical *P. aeruginosa* isolates, overall response rates were 81%, 39%, and 96% to the KIM-C1 cocktail, AZT, or the KIM-C1+AZT phage cocktail/aztreonam respectively in planktonic conditions. 80% of isolates grown as biofilms were likewise suppressed by KIM-C1+AZT (**Fig. 5B-ii**). Notably, some clinical *P. aeruginosa* isolates were not responsive either to KIM-C1 or AZT individually but were suppressed by KIM-C1+AZT (**Fig. 5B-iii**). These data indicate that a cocktail incorporating phages from the different CGs plus a synergic antibiotic was broadly effective against both planktonic and biofilm cultures.

We then extended this approach to generate additional cocktails (KIM-C2 and KIM-C3) of other phages encompassing multiple CGs. We found that these likewise achieved ≥96% coverage (**Fig. 5B-ii**). These data further demonstrate that the receptor-based complementarity grouping can inform the generation of effective phage-antibiotic cocktails with broad host range.

### Phage-antibiotic combination therapy is effective against multi-drug resistant clinical *S. aureus* isolates

We wondered whether this approach could be applicable to *S. aureus,* another AMR organism that often occurs in polymicrobial infections along with *P. aeruginosa*^10^. To this end, we examined a panel of five different *S. aureus* phages. For two of these (*S. aureus* bacteriophages K and Romulus), the receptors were known^51,52^ (**Fig. 6A-i**). We evaluated the Suppression Index for these phages against *S. aureus* strain 1203 **(Fig. 6A-ii, iii)**. Following re-exposure to these same phages, we used the resulting Resistance Index data to generate the phage exposure matrix shown in **Fig. 6B-i**. We observed that the phages targeting *S. aureus* fit into two major complementarity groups. This pattern was further supported by principal component analysis (PCA) (**Fig. 6B-ii**). As with our *P. aeruginosa* studies (**Fig. 2C**), the patterns of receptor usage did not always group with phylogenetic relationships (**Fig. 6B-iii**). We further identified characteristic patterns of antibiotic responses that exist for different *S. aureus* phage complementarity groups. In particular, vancomycin (VAN) was synergistic while rifampin (RIF) was antagonistic against phages belonging to both CGs (**Fig. 6C-i, ii**).

**Figure 6.**
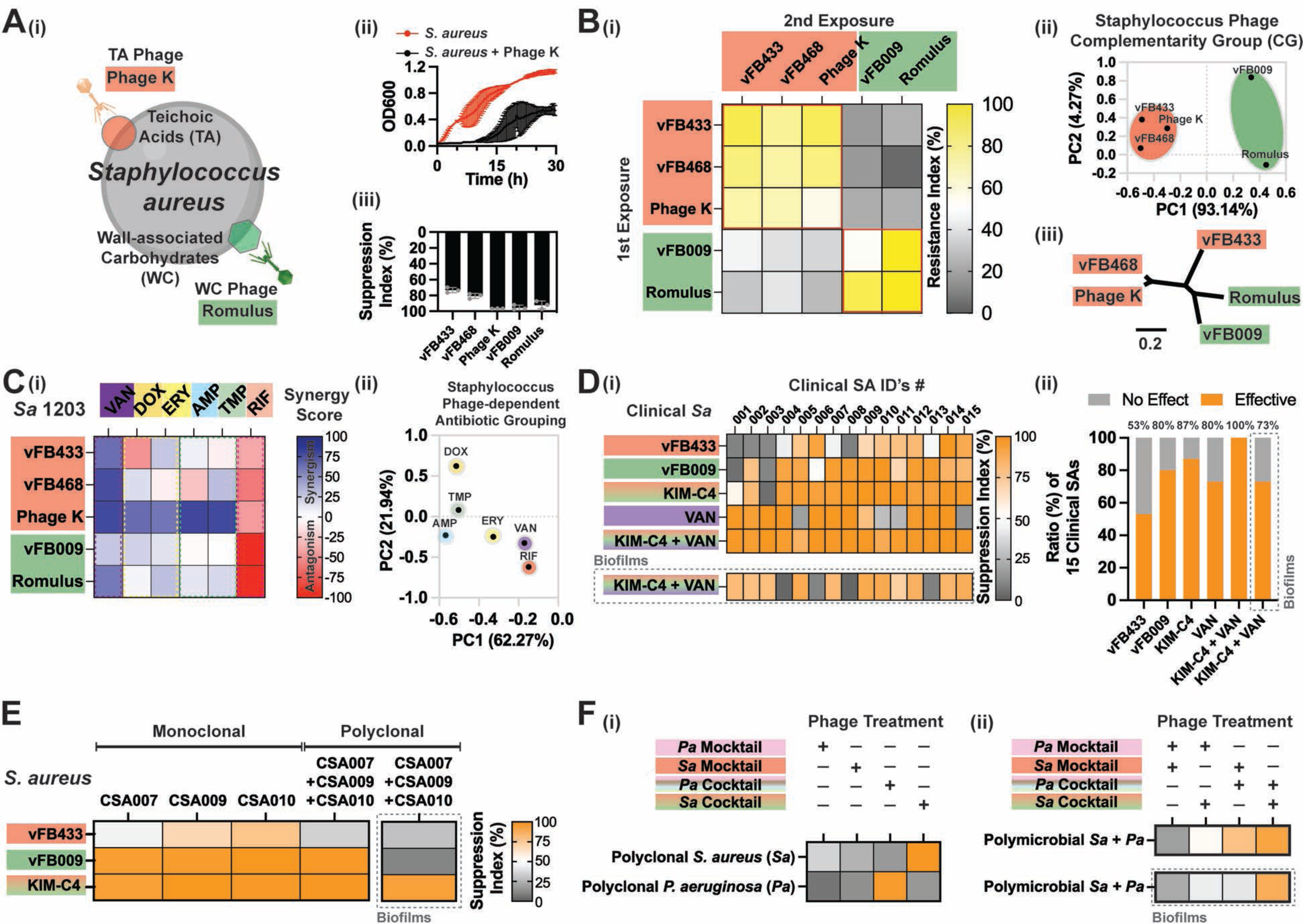
| Cocktails containing phages from three or more CG reliably kill multi-drug-resistant clinical isolates of *S. aureus* and also are effective for polymicrobial infections. (**A-i**) Schematic model description of two major surface receptors involved in phage uptake by *S. aureus.* (A-ii), Growth curves for *S. aureus* strain 1203 following treatment with phage K. **(A-iii),** Suppression Index for strain 1203 exposed to five different phages, each at 100 MOI. **(B-i),** Phage exposure matrix for strain 1203 initially exposed to one of five phages and then subsequently “re-exposed to each of the same five phages, all at MOI = 100. (**B-ii**), PCA analysis, and (**B-iii)** phylogenic dendrogram for these phages. **(C-i),** Synergic interaction matrix for 1203 exposed to combinations of phages and antibiotics simultaneously. VAN = vancomycin 1.0 μg/ml, DOX = doxycycline 0.05 μg/ml, ERY = erythromycin 1.0 μg/ml, AMP = ampicillin 0.1 μg/ml, TMP = trimethoprim 1.0 μg/ml, RIF = rifampin 0.03 μg/ml. **(C-ii),** PCA data for the data in **C-i**. (**D-i),** Suppression Index for the individual phages vFB433, vFB009 (100 MOI each), a cocktail (KIM-C4) containing both phages (100 of combined MOIs), vancomycin (VAN), or and phage/antibiotic cocktail (KIM-C4+VAN) grown in planktonic form or as biofilms. **(D-ii),** Effectiveness Ratio (%) of 15 clinical *S. aureus* isolates to the treatments in **D-i**. (**E-i),** Suppression Index for monoclonal versus polyclonal *S. aureus* isolates cultures (CSA007, CSA009, CSA010) treated individually with a cocktail (KIM-C4) or its two constituent phages (vFB433, vFB009). **F,** Suppression Index for polyclonal *S. aureus* (CSA007, CSA009, CSA010) or *P. aeruginosa* (CPA077, CPA086, CPA149) **(F-i)** or polymicrobial *S. aureus + P. aeruginosa* isolates (all six strains) **(F-ii)** cultures treated individually with cocktails of phages with the different CGs or “mocktails” of phages sharing the same CG as controls. *P. aeruginosa* mocktail (KIM-M1) includes DMS3vir, KOR_P1, and KOR_P2. *S. aureus* mocktail (KIM-M2) includes vFB433 and vFB468. *P. aeruginosa* cocktail (KIM-C1) includes Luz24, PAML-31-1 and OMKO1. *S. aureus* cocktail (KIM-C4) includes vFB433 and vFB009. While the polyclonal *S. aureus/P. aeruginosa* isolates cultures include three clinical isolates each, the polymicrobial *S. aureus* and *P. aeruginosa* cultures include all six isolates. All data shown here are from triplicate independent results.

Next, we considered whether cocktails informed by CGs had activity against *S. aureus* clinical isolates. To this end, we generated a cocktail called KIM-C4 including representatives of two different CGs using phages vFB433 and vFB009. We chose VAN as this antibiotic had widely synergistic interactions with all five phages in our set. We observed that most of a set of 15 clinical *S. aureus* isolates were occasionally susceptible to either these individual phages or vancomycin when used alone. However, the phage-antibiotic combination (KIM-C4+VAN) increased these responses substantially and had the broadest host range, including against biofilm cultures (**Fig. 6D-i, ii**). Polyclonal *S. aureus* cultures similarly had improved responses with the KIM-C4 cocktail (**Fig. 6E**).

Finally, we asked whether phage cocktails designed using CG principles were effective against polymicrobial cultures. To assess this, first we grew either three different *S. aureus* or three different *P. aeruginosa* isolates each at a 1:1:1 ratio. These polyclonal cultures were subsequently treated with either mocktails of phages sharing the same CG or cocktails of phages with different CGs. We observed that each polyclonal culture was specifically responsive to the species-specific cocktail (**Fig. 6F-i**). Next, we mixed, with the same ratio, a total of six clinical isolates (three different *S. aureus* and three different *P. aeruginosa* isolates in equal proportions). These polymicrobial cultures were subsequently treated with either “mocktails” of phages sharing the same CG or “cocktails” of phages with different CGs (**Fig. 6F-ii**). We observed that polymicrobial cultures simultaneously treated with both KIM-C1 (a *P. aeruginosa* phage cocktail) and KIM-C4 (a *S. aureus* phage cocktail) were most effectively suppressed, compared to the other conditions. Here, we highlight that we did not observe any antagonism between phages of each species, suggesting that mixtures of multiple species-targeting cocktails can be used for complex, polymicrobial infections.

Together, these data demonstrate that phage cocktails developed using CG principles are effective against both polyclonal and polymicrobial cultures that contain each representative Gram-negative and Gram-positive AMR pathogens.

## Discussion

Here, we report that by combining multiple phages from multiple, non-redundant receptor CGs we can create highly effective, broad-spectrum phage cocktails. Using this approach, we develop multiple cocktails active against large numbers of diverse clinical isolates of *P. aeruginosa* and *S. aureus,* including biofilm cultures and wound infections *in vivo*.

Along with broad host range, we report that cocktails including representative phages from multiple CGs can prevent the emergence of phage resistance, an important limitation of existing treatment protocols^11,13^. We propose that exposure to multiple phages that use different receptors may create orthogonal selection pressures that are difficult for bacteria to evade, as has been shown for at least one phage/antibiotic pairing^53^. Similar to the use of multiple anti-retroviral therapies with different mechanisms of action against HIV^54^, the use of at least three complementary phages may effectively suppress bacterial growth and prevent mutations leading to resistance.

Interestingly, we observed that CG patterns were distinct from the phylogenetic relationships derived from whole-genome sequences of the same phages. While phylogenetic relationships for particular genes might reveal other patterns, these results suggest that phage receptor usage is evolutionarily fluid, presumably due to the mosaicism of phage genome. In addition, it is noteworthy that some phages rely on multiple receptors, in agreement with previous reports^55^. Our results also point to previously unsuspected receptor interactions for some well-characterized phages, such as OMKO1^43^, which we find requires both OprM and the *P. aeruginosa* flagella for its infection. These findings suggest that phage-bacterial receptor interactions are likely dynamic and complex^8,41^, highlighting both the utility of a functional screening assay for phage-receptor interactions and the need for further study.

Particular phage CGs exhibit characteristic interactions with particular antibiotic classes. For example, phages belonging to the Type IV pili CG are consistently synergistic with β-lactam antibiotics and aminoglycosides but have idiosyncratic interactions with most other antibiotic classes. Interestingly, we observe that phage-antibiotic interactions are also strain/species-dependent. For example, rifampin would have always synergic interactions with *P. aeruginosa* phages, while having antagonistic interactions with *S. aureus* phages, presumably explained by different physiology of two organisms. While beyond the scope of this work, understanding the mechanisms underlying these patterns will be important for the future success of this approach. We envision that phage-antibiotic synograms^22^ could incorporate phage CG information to empower phage clinical treatment decisions, in the same way that physicians currently consult an antibiogram to inform treatment choices.

In addition to the conceptual advance of using phage cross-resistance to identify receptor CG and design effective cocktails, we also introduce toolkits to facilitate this approach. This includes systems for quantifying phage suppression and resistance and a robust in vitro/in vivo model for evaluating cocktail activity against polymicrobial infections. While several of these elements have antecedents in the literature^22,56,57^, their incorporation into an integrated system, will facilitate further development of phage-antibiotic cocktails for these and other pathogens.

Future studies may build on these findings in other ways. In particular, it may be possible to further expand the existing CG matrices reported here to include additional characterized or unknown phages. In addition, by expanding the CG matrices, we could generate additional phage cocktails against *P. aeruginosa* and *S. aureus*. We used *P. aeruginosa* and *S. aureus* for these studies because of the urgent medical need for new treatments against these deadly pathogens. However, it should be possible to generate phage cocktails against other Gram-positive and Gram-negative pathogens as well, using the same technique, analytic tools, and principles described here. The potential of this therapeutic approach could be further extended by incorporating molecules that degrade biofilm matrix such as holins^58^, given that we observed the reduced effectiveness of cocktails within biofilms. Our strategy further suggests the potential to be expanded to other systems like fungal infections using mycophages, a promising avenue for future research.

In conclusion, these studies establish a blueprint for the rational design of broad-spectrum phage cocktails that may overcome the serious medical problem of highly resistant bacterial infections. We hope that the approaches developed here will enable future off-the-shelf phage-based therapeutics and thereby unlock the clinical and commercial potential of phage therapy.

## Acknowledgments

We thank Dr. Zemer Gitai (Princeton University), Dr. Bonnie Bassler (Princeton University), Dr. Bob Hancock (University of British Columbia), Dr. George O’Toole (Dartmouth College), Dr. Pat Secor (University of Montana), Dr. Rob Lavigne (Katholieke Universiteit Leuven) and the members of Felix Biotechlogy, Inc (San Francisco, CA) for gifting us bacterial stains and bacteriophages (See the details for Supplementary Information). We also thank Drs. Gina Suh, Taia Wang, Manish Butte, Markus Covert, and Trung Pham for their valuable feedback, Tejas. Dharmaraj and Yung-Hao for assistance with *in vivo* animal studies; and all members of the Bollyky Laboratory for insightful discussions. This work was supported by NIH R01HL148184-01, NIH R01AI12492093, NIH R01DC019965, the Cystic Fibrosis Foundation, and the Emerson Collective (to P.L.B). The Severance Alumni Moon Scholarship Foundation (to M.K.K). A scholarship from Sue and Thomas Merigan (to A.E). The funders had no role in the study design, data collection, analysis, the decision to publish, or the preparation of the manuscript.

## Author Contributions

M.K.K. conceptualized the main idea of this research project, designed experiment methodology, and performed the majority of the experiments. P.L.B. provided feedback on the results. M.K.K. and P.L.B. formally analyzed the data and wrote the original draft. Q.C. and A.E. performed *in vivo* experiments and the relevant data analyses. N.P. performed the TEM imaging. R.C.M., N.B., and E.B.B., and C. E. M. collected and provided clinical isolates and bacteriophages. All authors contributed to the revision and editing of the paper.

## STAR Methods

### CONTACT FOR REAGENT AND RESOURCE SHARING

Further information and requests for reagents may be directed to and will be fulfilled by the corresponding authors, (M.K.K) and (P.L.B).

## METHOD DETAILS

### Bacterial strains and overnight growth conditions

The strains used are listed in Supplementary Table 3. *P. aeruginosa* strains PA14 were used as a representative *P. aeruginosa* strain for most experiments shown in Fig. 1, 2, 3, 4, and 5 and their corresponding supplementary figures unless stated otherwise. For experiments shown in Fig. 4, 5, and 6, clinical *P. aeruginosa* strains were used. For the experiment shown in Fig. 6, *S. aureus* strain 1203 was used as a representative *S. aureus* strain, and 15 clinical isolate strains of *S. aureus* were also used. For growth conditions, lysogeny broth (LB) was used as media and if applicable, appropriate antibiotics were added. LB agar contained LB with 18 g/L agar, and LB top agar included LB with 3.0 g/L agar, 50 mM CaCl_2,_ and 1 mM MgSO_4_.

Collection of clinical *P. aeruginosa* and *S. aureus* isolates from patients *P. aeruginosa* and *S. aureus* were isolated from various patient sources including sputum, BAL, sinus, and wounds in the Stanford Healthcare Clinical Microbiology Laboratory (SHCML), from June 2020 to May 2023. After 153 *P. aeruginosa* and 15 *S. aureus* clinical isolates were collected, all strains were de-identified. The clinical samples were cultured on routine media (blood, chocolate, MacConkey, and CNA agar). Pathogens were identified by MALDI-TOF (Bruker Biotyper). These isolates were given a unique identifier and stored at –80 °C in 40% glycerol until further use.

### Bacterial growth curves for phage suppression assays

Cultures from frozen stocks were incubated at 250 rpm shaking at 37 °C overnight. Overnight cultures of bacterial strains were back-diluted 1:200, and re-grown for 2-3 hours (to OD600 ∼ 0.1-0.2). Then, the re-grown culture was diluted 1:100 (OD600 ∼ 0.001-0.002) into wells on a 96-well plate with LB to a total volume of 150 μL with pre-mixed phage (phage cocktails) to reach the target MOI. Depending on the experiment condition, an appropriate antibiotic was also added to each well to reach the target concentration. The final cultures on a 96-well plate were then incubated at 37 °C with shaking for 30 hours, and the OD600 curve was read every 20 min using an automated spectrophotometer (Biotek microplate reader). All the growth curves were subtracted from the background curve (LB media with phage buffer (1 mM MgSO_4_, 4 mM CaCl_2_, 50 mM Tris-HCl, pH=7.8, 6g/L NaCl and 1g/L gelatin)). Using the built-in GraphPad Prism Version 7.0 software package, the Suppression Index (%) was calculated as the area under the curve (AUC) for the non-phage-treated condition minus the phage-treated condition divided by the AUC for the non-treated condition.

### Bacterial growth curves for phage resistance assays

Phage-resistant isolates were obtained as follows. First, the strains after 30 hours of exposure to the 1st phage (or cocktails) were plated on the LB agar, where two colonies were randomly selected in the most dilute spots. Each picked colony was re-streaked onto sterile LB plates and incubated overnight at 37 °C. Each re-streaked isolate was picked and then suspended in sterile LB and incubated overnight at 37 °C with 250 rpm shaking. 500 μL samples of each bacterial culture were combined with 500 μL of sterile 60% glycerol solution and frozen at −80 °C. These stocks were used for overnight cultures. The growth conditions were the same as described above in the suppression assay. The Resistance Index (%) was calculated as the AUC of the phage-rechallenged condition divided by the AUC of the non-rechallenged condition after 15 hours.

### Phage strains, purification, quantification, and plague assay

The bacteriophages and their host strains for phage propagation are listed in Supplementary Table 4. For phage amplification, host bacteria strains at the mid-log phase were infected with stocks of phage and then cultured in 5 mL of LB top agar solution, which was subsequently poured into a sterile LB agar plate and incubated overnight at 37 °C. Phages were collected by adding 5 ml of phage buffer into the agar and collecting the top agar overlays. After collection of the top agar pieces into a 50 ml tube, any agar residuals were spun down by centrifuging them at 10,000 × g for 30 min. The supernatants in the tube were collected and filtered via a 0.45 mm PES syringe. The collected phages were proceeded for further quantification. Phages were precipitated from the collected supernatant by adding 0.5 M NaCl and 4 % (w/v) polyethylene glycol (PEG) 8000 (Milipore) and then were incubated overnight at 4 °C. Phages were pelleted by centrifugation at 13,000 × g for 20 min, and the pellet was then suspended in sterile TE (Tris, EDTA) buffer (pH 8.0). The suspension was centrifuged for 15,000 × g for 20 min at 4 °C, and the supernatant was subjected to another round of PEG precipitation. The purified phage pellets were suspended in sterile PBS and dialyzed in 10 kDa molecular weight cut-off tubing (Fisher Scientific) against PBS, diluted to appropriate concentrations in sterile phage buffer, and filter-sterilized. For phage quantification, a phage titer in the filtered phage suspensions was determined as follows. Tenfold serial dilutions of phage suspensions were prepared in phage buffer, ranging from 10^−1^ to 10^−12^. From each dilution, 10 μL was spotted on the host bacterial lawn on LB agar and incubated overnight at 37 °C. After incubation, plaques were counted at the spots at which the phage suspensions were added. The final PFU (plaque forming unit) was made based on the two independent counting. For plaque assay, 10 μL of the phage was spotted on the host bacterial lawn and incubated overnight at 37 °C. After incubation, pictures of phage plaques were taken from three independent plates, as shown as representative images in the Fig. 1.

### Isolation of wild phage KOR_P1 and KOR_P2

The phages were isolated from the Codiga Resource Recovery Center (CR2C) at Stanford (692 Pampas Lane, Stanford, CA 94305, United States). Wastewater from the plant was centrifuged and then the supernatant was filtered (pore size = 0.22 μm). PAO1Δpf4Δpf6 was used as a host and then mixed with the sewage and grown on an agar overlay overnight at 37 °C. Clear individual plaques were selected and suspended in the phage buffer. Using this phage-containing buffer, this process was repeated three times, and phages KOR_P1 and KOR_P2 were plaque-purified and then quantified for their PFU.

### Isolation of phage genomic DNA

Phage genomic DNA from KOR_P1 and KOR_P2 was extracted by the following method using 0.5 ml of lysate (10^10^ pfu/ml). To remove any residual bacterial DNA and RNA present in the lysate, 0.48 ml of the filter-sterilized lysate is incubated with 1 μL DNase I (1 U/μL) and 1 μL RNase A (10 mg/mL) for 90 min at 37 °C without shaking. Thereafter, 20 μL of 0.5M EDTA (final concentration 20 mM) was added to inactivate DNase I and RNase A. To digest the phage protein capsid, 1.25 μL Proteinase K (20 mg/mL) is then added and incubated for 90 min at 55 °C without shaking. The digested phage DNA was then purified with the DNeasy Blood & Tissue Kit (Qiagen) with the modified manufacturer manual^59^.

### Transmission Electron Microscopy

Images of high-titer phage lysates (5 µl) were collected as follows. The phage lysates were absorbed onto glow discharged forever and carbon-coated copper grids (300 nm mesh size, Electron Microscopy Sciences) at room temperature for 5 min, negatively stained with 1% (w/v) uranyl acetate at room temperature for 1 min and imaged with a JEM-1400 transmission electron microscope (Jeol) operated at 120 kV. The images were taken with a GATAN Multiscan 791 CCD camera.

### Construction of phylogenetic trees for bacteriophages

Sequences of phage DNA were obtained from the database found in NCBI Nucleotide. The sequences of phage TIVP-H6, vFB009, vFB433, and vFB468 were provided from the original source (Felix Biotechnology, Inc). Using the Tree Builder in the Geneious Prime software with the default settings, phylogenetic trees were generated and the genetic distances between phages were obtained.

### Whole Genome Sequencing (WGS), Genome Assembly, Annotation, and Comparison

Libraries of the selected bacterial DNA samples (input DNA 0.2 ng/μl) were prepared using the Illumina Nextera XT DNA Sample Preparation Kit following the manufacturer’s instructions. Sequencing of DNA (paired– end 2 × 150 high output) was carried out using the Illumina NovaSeq PE150 platform. After adapter trimming, sequence reads were assembled using the Geneious Prime v2023.1.2. All individual genome assemblies were annotated and assessed through mapping to the reference *P. aeruginosa* PA14 bacterial genome NC_008463 with default settings. Single nucleotide polymorphism (SNP) detection was conducted using the Geneious Prime software with the default settings.

### Suppression assays using knockouts library of *P. aeruginosa* PA14

The knockout mutants used for the suppression assays included 12 strains, which are listed in the Supplementary Table 1 and differed in the knockout of a gene for a surface-expressed protein for the phage receptors: type IV pili genes (*pilA, pilB, pilC, pilE, pilW, pilTU, pilX,* and *pilY1*), flagella genes (*filC* and *flaK*), a Mex system protein gene (*oprM*), and a OSA/LPS synthesis gene (*wapR*). Suppression assays are the same as mentioned above. The suppression ability of a phage on the test knockout strain was compared to its ability on the phage-sensitive host (PA14).

### Motility (twitching and swimming) assays

Following an established protocol^60^, bacterial cultures were grown overnight and diluted and regrown to OD = 0.1-0.2. Using a pipette tip, the bacterial suspension was transferred by a puncture into the agar plates (containing a 1% tryptone medium with different concentrations of agar (0.3% for swimming motility and 2% for twitching motility)). The plates were then incubated at 37 °C for 24 hours (swimming motility) and 48 hours (twitching motility). After the incubation, the growth zones were measured. For swimming motility, the measurement was made directly. For twitching motility, the agar layer was removed and the bottom of the plate was stained with 5 ml of a 0.01% solution of crystal violet (Sigma-Aldrich) for 10 min and washed with DI water. Each experiment was performed for at least 5 repetitions.

### Colony biofilm morphology assay

Following an established protocol^61^, 1 μL of overnight PA14 cultures was spotted onto the agar plates (containing a 1% TSB medium fortified with 40 mg/L Congo red and 20 mg/L Coomassie brilliant blue dyes and solidified with 1% agar). Biofilms were grown at 25 °C, and images were acquired after 100 hours, using a Leica M80 Stereo Microscope mounted with a Nikon 170 camera at 7.5 × zoom.

### Hitchcock and Brown LPS Preparation, LPS SDS-PAGE, and LPS modified silver staining

LPS was prepared following the Hitchcock and Brown LPS Preparation method^62^. 3 μL of LPS solutions were treated for 5 min at 100 °C after mixing with 17 μL water and 20 μL sample buffer. The 40 μL mixture was transferred into each well of a 12% discontinuous PAGE mini-slab Tris-glycine-polyacrylamide gels and poured with the SDS-running buffer (0.25 M Tris, 1.92 M glycine, 1 % w/v SDS)) in a vertical gel electrophoresis system, and then ran at 200 V for 50 min. The fractionated LPS-SDS-PAGE-gels were first oxidized with 0.7% periodic acid in 40% ethanol-5% acetic acid at 25 °C for 30 min without any prior fixation. The gels were washed twice with DI water and followed the protocol of the Pierce™ Silver Stain Kit (Thermo Scientific). The final images were taken by a Bio-Rad ChemiDoc MP imaging system.

### Surface biofilm imaging assays

Following the previous protocol for the biofilm imaging^47,63^, the overnight cultures from those strains that harbor a gene that constitutively expresses fluorescent proteins were regrown and diluted to OD600 ∼ 0.001 and 100 μL of the diluted cultures were added to wells of 96-well plates with no. 1.5 coverslip bottoms (MatTek). The cells were allowed to attach for 10 min, after which the wells were washed twice with fresh LB medium, and, subsequently, 100 μL of fresh LB medium was added and grown at 25 °C. Images were acquired with a Yokogawa CSU-X1 confocal spinning disk unit mounted on a Nikon Ti-E inverted microscope, using a 60× water objective with a numerical aperture of 1.2, the 405/488/561/642 multichroic laser, and a Photometrics Prime 95B camera. All experimental images in this work were acquired and rendered using Nikon Element software. Each biofilm covering a surface area of 109 × 109 μm^2^ was imaged at six different regions for each well in at least n = 3 independent experiments. Each image was then segmented in the z-plane and assessed independently to count the cells in the biofilms using the previously published custom code^64^.

### Modified microtiter dish biofilm formation assay

Following an established protocol^65^ with modifications, strains that do not harbor a gene that constitutively expresses fluorescent proteins (e.g., clinical isolated strains) were cultured overnight, regrown, and diluted to OD600 ∼ 0.001. 15 μL of the diluted cultures were then added to each 96-well plate with 135 μL of LB media containing phage cocktails, as applicable. The plate was then covered with a pegylated lid and sealed with the sides of the plate with a permeable membrane to prevent evaporation and incubated for 48 hours at 25 °C with shaking of 100 rpm. After 48 hours, the biofilm-attached pegylated lid was removed and placed into the fresh plate that contains sterile PBS. Then, the plate was shaken for 1 min to wash out the un-attached bacteria from the lid. After this process, the lid was placed on a fresh plate with 200 μL of sterile LB media in each well and sonicated for 1 h to dislodge biofilm cells from the lid by the vibrations created by the sonicator. After removing the lid from the plate, biofilm cells in the plate were amplified for 24 hours and then OD600 was measured to quantify the cells. The Suppression Index (%) was calculated as the OD600 for the non-treated condition minus the treated condition divided by the OD600 for the non-treated condition.

### Antibiotic-phage synergic assays

*P. aeruginosa* strain PA14 and *S. aureus* strain 1203 were grown overnight at 37 °C as described above. The cultures were back-diluted 1:200, and re-grown for 2-3 hours (to OD600 ∼ 0.1-0.2). Then, the re-grown cultures were diluted 1:100 (OD600 ∼ 0.001-0.002) into wells on a 96-well plate with LB to a total volume of 150 μL with/without a phage and/or antibiotics to reach the target MOIs and serial concentrations. The plates were incubated at 37 °C with shaking for 30 hours, the OD600 growth curves were obtained, and the bacterial levels (a.u., defined as the AUCs of OD600 for 30 hours) were obtained from triplicate experiments for each concentration of either an antibiotic or a phage individually and both.

Synergy scores were obtained for the antibiotic-phage interaction using the Highest Single Agent model^66,67^. In brief, bacterial cells grow under specific concentrations involving either an antibiotic or a phage. The impact of the more effective agent (by either an antibiotic or a phage) was defined as the expected effect of the combination of both an antibiotic and a phage. If the observed growth suppression in combination was equal to the expected value, each of the antibiotic and a phage was deemed to be independent. If the growth suppression was more or less than expected, the antibiotic and a phage were deemed to be synergistic or antagonistic, respectively. When an antibiotic and a phage were combined, the observed effect of the combination could be high or low, compared to what was expected given the individual effects of a given agent. The synergic scores were measured by comparing the growth suppression to an antibiotic and a phage and their combinations.

### *In vivo* murine full-thickness wound infection model

All male C57BL/6J mice for the wound infection experiments were purchased from The Jackson Laboratory (Bar Harbor). Mice were maintained under specific pathogen-free conditions, with free access to food and water, in the vivarium at Stanford University. All experiments and animal use procedures were approved by the Institutional Animal Care and Use Committee (IACUC) at the School of Medicine at Stanford University. The study design was adapted from the previous work^48^. In brief, 6–8-week-old male mice were anesthetized using 3% isoflurane. The dorsum of mice was shaved using a hair clipper and depilated using Nair hair removal cream (Church and Dwight). The shaved area was cleaned with betadine (Purdue Frederick Company) and alcohol swabs. Mice received 0.1-0.5 mg/kg slow-release buprenorphine (ZooPharm) subcutaneously before wounding. Bilateral dorsal full-thickness wounds were created using 6-mm biopsy punches (Integra). The wounds were covered with Tegaderm (3M). PAO1, PA14, and CPA 050 were grown as described above and diluted and mixed to 1 x 10^7^ CFU/mL in a 1:1:1 ratio. Mice received 5 x 10^4^ CFU/wound bacteria via injection into each wound under the Tegaderm patch. All mice received the first treatment at two hours post-infection and the second treatment at 24 hours post-infection. A phage cocktail (KIM-C1) was prepared on the same day of the treatment in MOI = 200 in a 1:1:1 ratio for each individual phage. Control mice were treated with phage buffer. Mice were weighed daily and provided with Supplical Nutritional Supplement Gel (Henry Schein Animal Health). Three days post-infection, mice were euthanized by CO_2_ chamber and cervical dislocation and the wound bed was excised, homogenized, and plated for CFU analysis.

## QUANTIFICATION AND STATISTICAL ANALYSIS

### Statistical Analysis

All experiments were performed with a minimum of three replicates, and these values were used to plot mean ± standard deviation. Statistical analysis was performed using GraphPad Prism Version 7.0 software package, and data were analyzed using unpaired t-tests, Mann-Whitney test, ordinary one-way analysis of variance (ANOVA), and multiple comparison test to determine the significance of results. Results were taken as significantly different by a p-value of < 0.05 unless otherwise stated.

## DATA AND SOFTWARE AVAILABILITY

### Data availability

All materials and experimental data that support the findings of this study are available without restriction by request from the corresponding authors. Source data are provided in this paper.

### Code availability

All custom codes used to process biofilms are available in the previously published paper^64^.

## SUPPLEMENTAL INFORMATION

Supplemental Information includes five figures and four tables

