## Supplementary files for "A Blueprint for Broadly Effective Bacteriophage Therapy Against Bacterial Infections"

Minyoung Kevin Kim, Qingquan Chen, Arne Echterhof, Robert C. McBride, Nina Pennetzdorfer, Niaz Banaei, Elizabeth B. Burgener, Carlos E. Milla, and Paul L. Bollyky

**This file includes:**

Supplementary Figures S1-S5

Supplementary Tables S1-S4

**Supplementary Figures:**

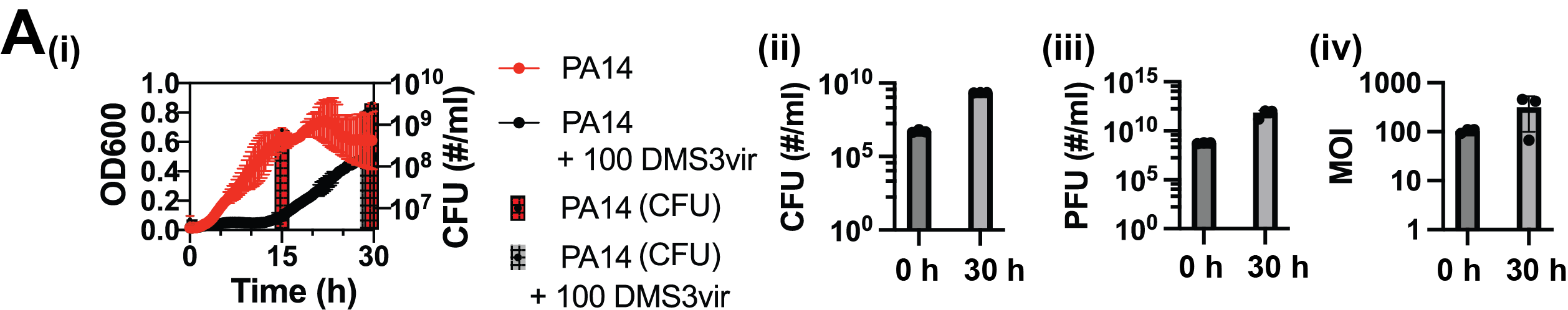

**Supplementary Figure 1 | Repeated exposure to lytic bacteriophages leads *P. aeruginosa* to develop constitutive, high-level resistance to those phages. (A-i**), Paired growth curves and colony forming unit (CFU) data for PA14 treated with or without 100 MOI by DMS3vir phage. CFU was measured at 0 h, 15 h, and 30 h whereas OD600 was measured continuously. (**A-ii**), CFU of PA14 in the presence of DMS3vir phage at 0 and 30 h. (**A-iii**), plaque-forming unit (PFU) of DMS3vir phage at 0 h and 30 h. (**A-iv**), MOI of DMS3vir phage was measured at 0 and 30 h.

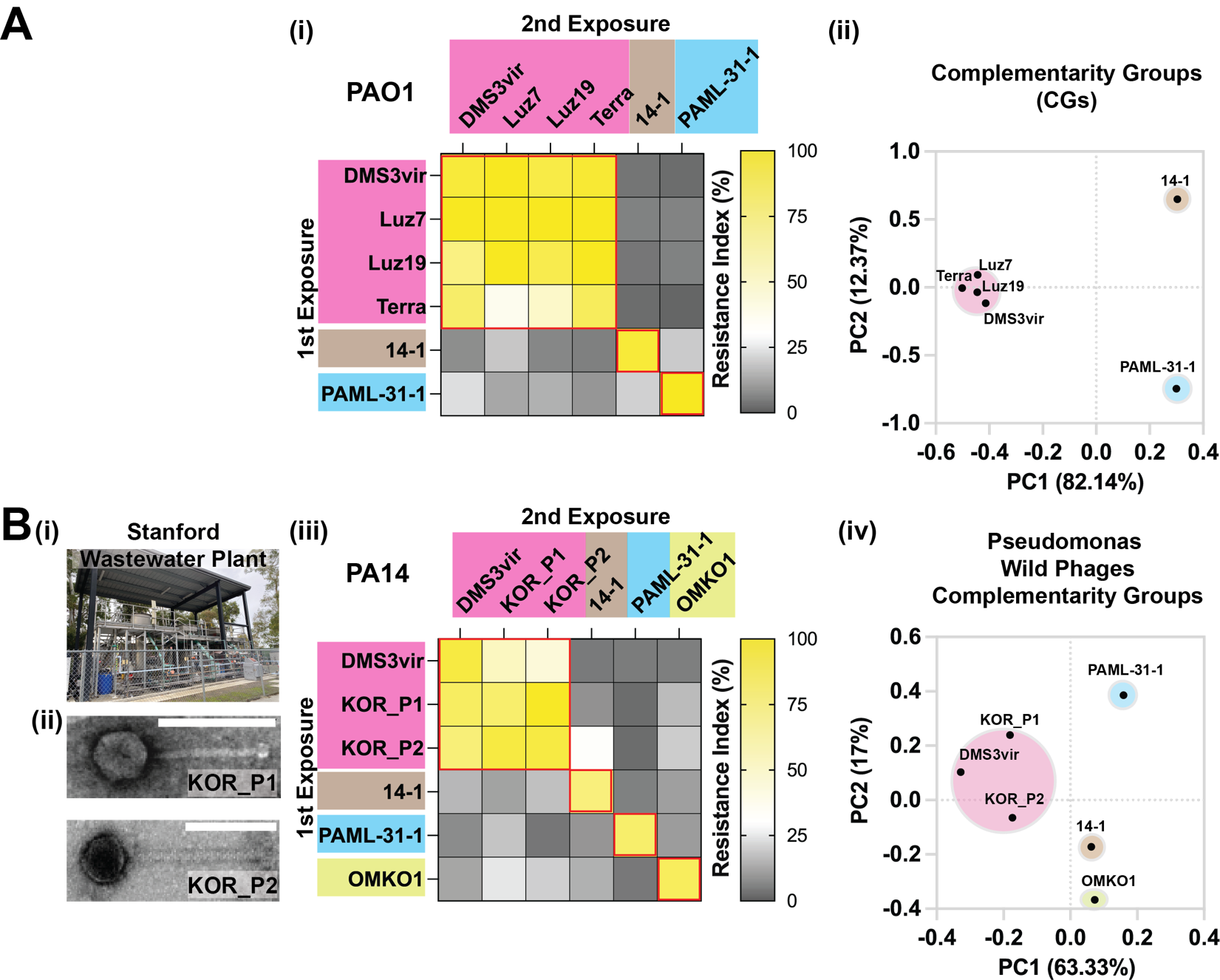

**Supplementary Figure 2 | The same CGs observed for *P. aeruginosa* strain PA14 can be observed for *P. aeruginosa* strain PAO1 and also accommodate novel phages. (A-i),** Resistance Index matrix for phages targeting *P. aeruginosa* strain PAO1 demonstrating the presence of the same complementarity groups seen for PA14. PAO1 was initially exposed to one of a set of six phages and then subsequently “re-exposed” to each of the same set of six phages, all with MOI = 100. Data for the Resistance Indices in the chart denote the means from triplicate experiments. **(A-ii),** PCA analysis of the chart description of (a-i). **(B-i),** The Stanford Wastewater Plant from which we isolated two previously “uncharacterized” phages. **(B-ii),** Transmission electron microscopy (TEM) images of the two phages called KOR_P1 (top) and KOR_P2 (bottom) isolated at the plant. Images are based on triplicate independent replicates. Scale bars, 100 μm. **(B-iii),** Resistance Index matrix for the four major complementarity groups including phages KOR_P1 and KOR_P2. Each row shows the Resistance Index (%) from each designated phage-exposed PA14 that was re-challenged by six different phages with MOI = 100. Data represent the means from triplicate experiments. **(B-iv),** PCA analysis of the chart description of **(B-iii)**.

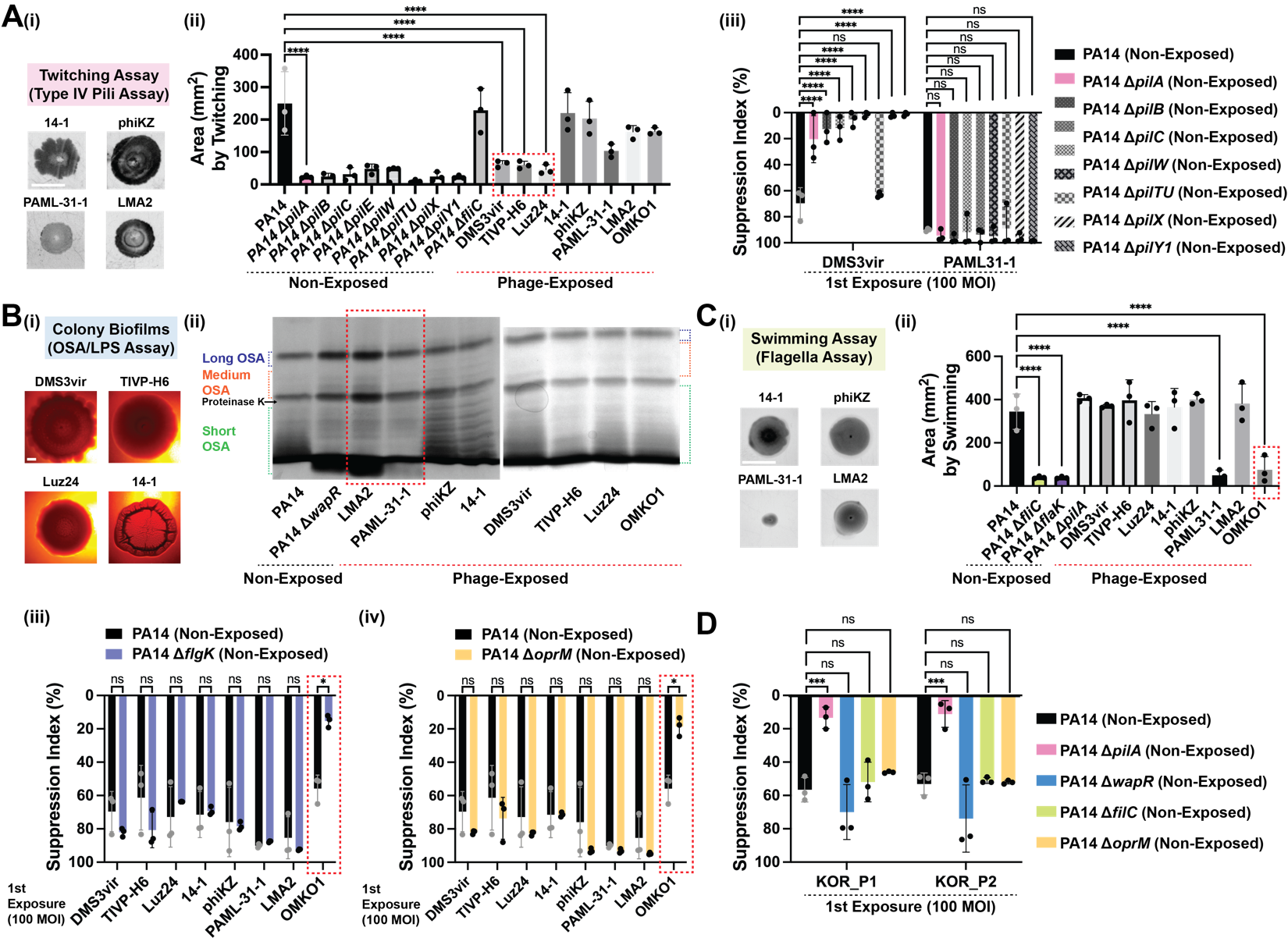

**Supplementary Figure 3 | CGs of phages target the same *P. aeruginosa* receptor structures. (A-i),** Representative images of twitching motility assay by 14-1-, phiKZ-, PAML-31-1-, and LMA2- exposed *P. aeruginosa* strain PA14. **(A-ii),** Surface area (mm^2^) covered by twitching motility of wild-type, the designated mutants, and phage-exposed PA14 on 2.0% agar plates. **(A-iii),** Suppression Index measured for PA14 and PA14 Type IV mutants (PA14Δ*pilA*, PA14Δ*pilB*, PA14Δ*pilC*, PA14Δ*pilE*, PA14Δ*pilX*, PA14Δ*pilTU*, PA14Δ*pilX*, and PA14Δ*pilY1*) by DMS3vir and PAML31-1, each at MOI = 100. **(B-i),** Colony biofilm phenotypes of the DMS3vir-, TIVP-H6-, Luz24-, and 14-1-exposed PA14 strains on Congo red agar medium after 100 h of growth (scale bar = 2 mm). **(B-ii),** Characterization of the phenotype of oligosaccharide antigen (OSA)-containing LPS by silver-stained SDS-PAGE gel for PA14, PA14Δwa*pR*, and the designated phage-exposed PA14. **(C-i),** Representative images of swimming motility assay by 14-1-, phiKZ-, PAML-31-1-, and LMA2- exposed *P. aeruginosa* strain PA14. **(C-ii),** Surface area (mm^2^) covered by swimming motility of wild-type, the designated mutants, and phage-exposed *P. aeruginosa* on 0.3 % agar plates. **(C-iii),** Suppression Index was measured for PA14 (black) and the PA14 flagella mutants, PA14Δ*flaK* (purple) treated with eight different phages, each with MOI = 100. **(C-iv),** Suppression Index was measured for PA14 (black) and the PA14*ΔoprM* (yellow) by eight different types of phages, each with MOI = 100. **D,** Suppression Index of KOR_P1 and KOR_P2, each at MOI = 100, measured following treatment of PA14, PA14Δ*pilA,* PA14Δ*wapR*, PA14Δ*fliC*, and PA14Δ*oprM*. Error bars in these figures denote standard deviations from triplicate experiments. *P < 0.05, **P<0.005 and ***P < 0.0005 for multiple unpaired t-tests. Images are based on a minimum of triplicate independent replicates. Scale bars = 10 mm.

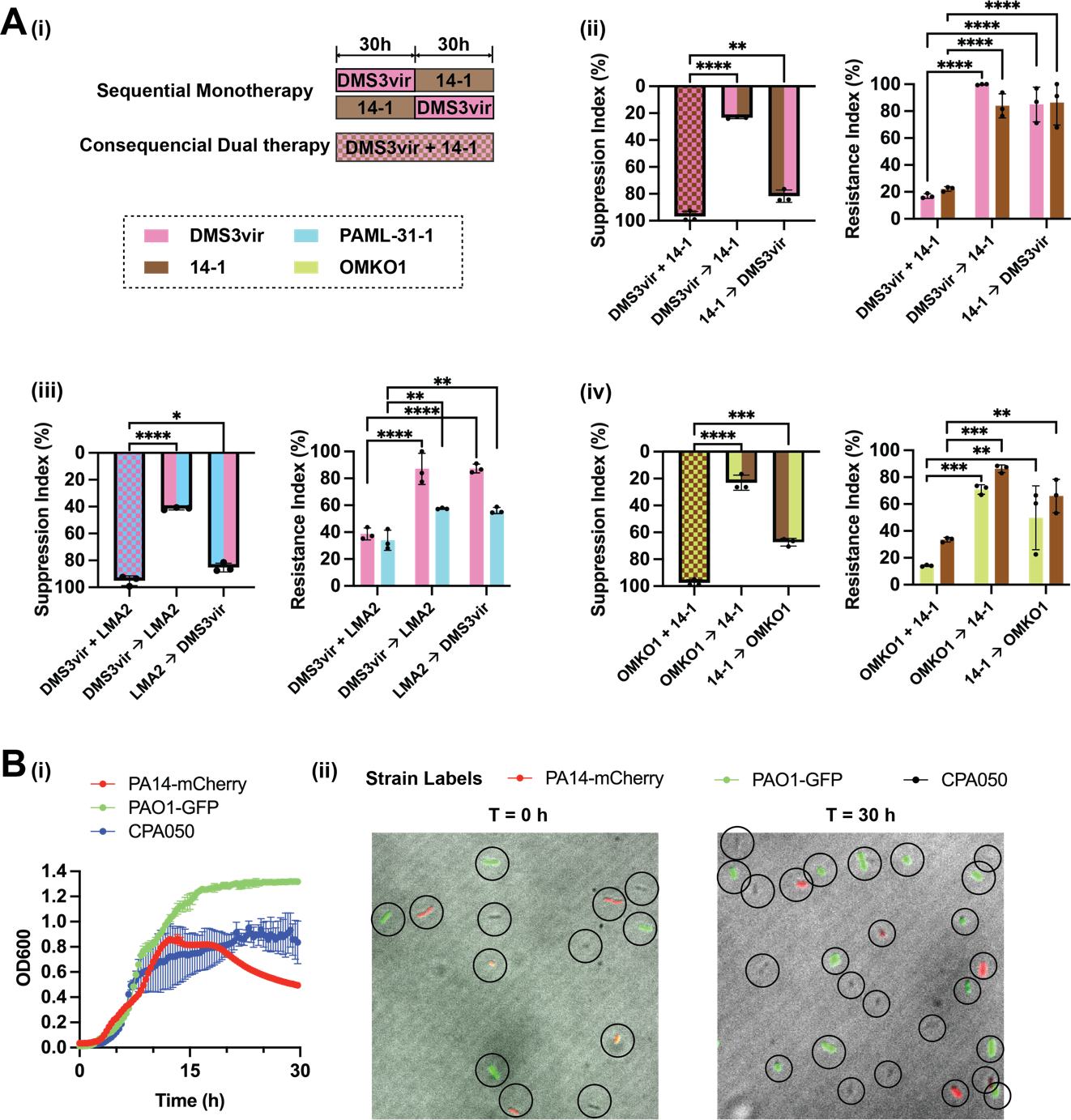

**Supplementary Figure 4 | Cocktails containing phages from different CGs effectively kill bacteria without the emergence of phage resistance. (A-i),** Schematic of sequential monotherapy versus consecutive dual therapy over 60 hours. For sequential monotherapy, treatment is initially with one phage (DMS3vir, pink) with MOI = 100 followed by a phage from a different CG (14-1, brown) with MOI = 100 after 30 hours. For consecutive dual therapy, two phages from different CG (DMS3vir + 14-1) each at MOI = 100 are added at both 0 and 30 hours. **(A-ii),** Suppression Index for consecutive dual therapy (DMS3vir, 14-1) versus two sequential monotherapies. Resistance Index from PA14 that was isolated after 60 hours of exposure and subsequently re-challenged with each constituent of the cocktail pattern as shown, each with MOI = 100. **(A-iii),** Suppression Index for consecutive dual therapy (DMS3vir, LMA2) versus two sequential monotherapies. Resistance Index from PA14 that was isolated after 60 hours of exposure and subsequently re-challenged with each constituent of the cocktail pattern as shown, each with MOI = 100. **(A-iv),** Suppression Index for consecutive dual therapy (OMKO1, 14-1) versus two sequential monotherapies. Resistance Index from PA14 that was isolated after 60 hours of exposure and subsequently re-challenged with each constituent of the cocktail pattern as shown, each with MOI = 100. **(B-i**), Growth curves for PA14-mCherry, PAO1-GFP, and CPA050. Error bars in this figure denote standard deviations from triplicate experiments. **(B-ii**), Representative images from triplicate experiments of individually discernable fluorescently labeled cells (Red: PA14, Green: PAO1, and Black: CPA050), which were sampled from mixed cultures (T = 0 h, left and T = 30 h, right).

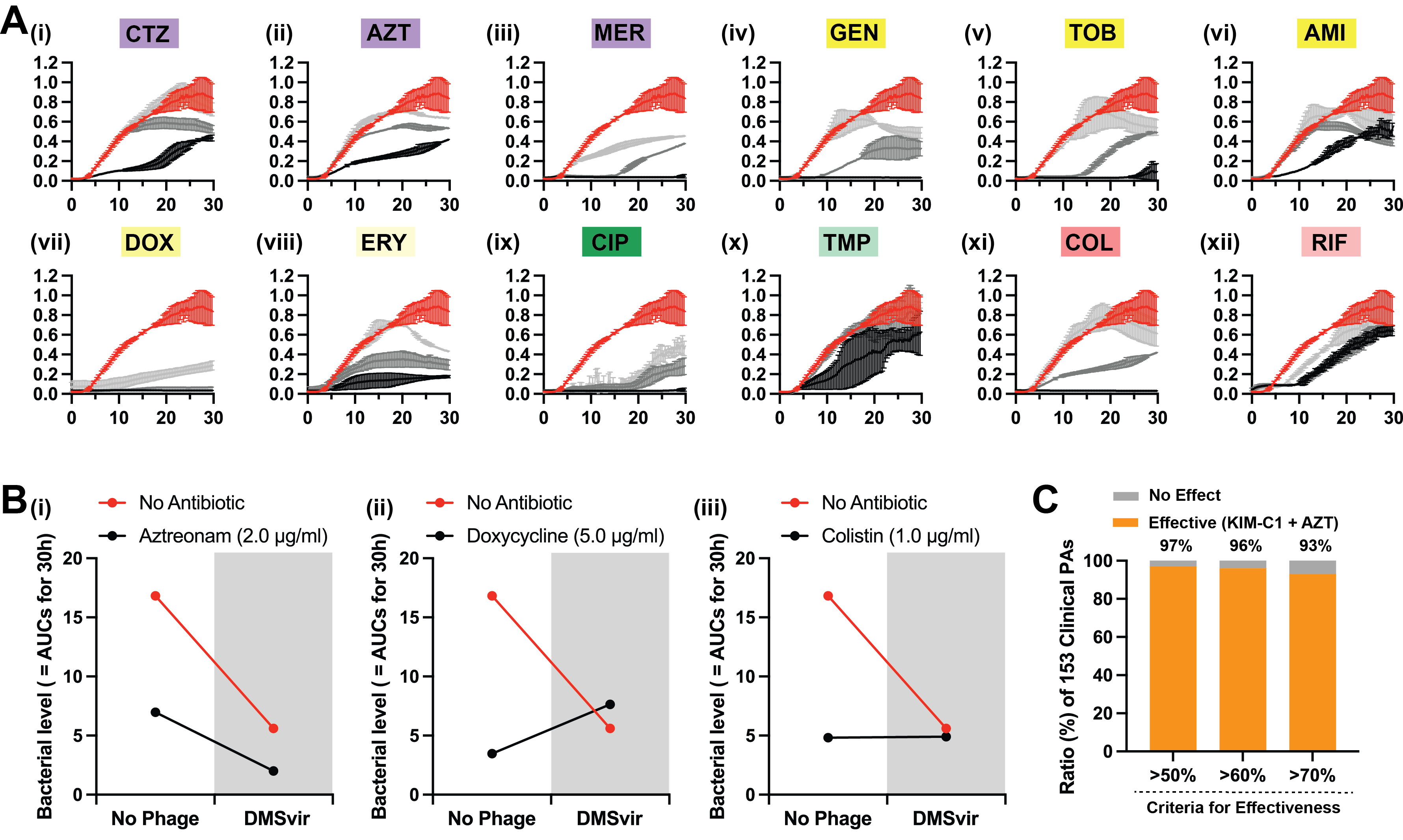

**Supplementary Figure 5 | Predictable, class-dependent interactions of conventional antibiotics with phage CG. (a)** Growth curves (OD600) from wild-type PA14 grown under different sub-MICs of 12 conventional antibiotics used in these experiments: Red curves = no antibiotics in all growth curves. **(A-i),** Aztreonam (AZT) responses. Black, dark grey, and grey = 2, 1, 0.5 μg/ml AZT, respectively. **(A-ii),** Meropenem (MER) responses. Black, dark grey, and grey = 0.5, 0.25, 0.125 μg/ml MER, respectively. **(A-iii),** Gentamycin (GEN) responses. Black, dark grey, and grey = 2.0, 1.0, 0.5 μg/ml GEN, respectively. **(A-iv),** Tobramycin (TOB) responses. Black, dark grey, and grey = 0.5, 0.25, 0.125 μg/ml TOB, respectively. **(A-v),** Amikacin (AMI) responses. Black, dark grey, and grey = 1.5, 0.5, 0.25 μg/ml AMI, respectively. **(A-vi),** Doxycycline (DOX) responses. Black, dark grey, and grey = 20.0, 10.0, 5.0 μg/ml DOX, respectively. **(A-vii),** Erythromycin (ERY) responses. Black, dark grey, and grey = 40.0, 20.0, 10.0 μg/ml ERY, respectively. **(A-viii),** Ceftazidime (CTZ) responses. Black, dark grey, and grey = 1, 0.5, 0.25 μg/ml CTZ, respectively. **(A-ix),** Ciprofloxacin (CIP) responses. Black = 2.0, 1.0, and 0.5 μg/ml CIP, respectively. **(A-x),** Trimethoprim (TMP) responses. Black = 200, 100 and 50 μg/ml TMP, respectively. **(A-xi),** Colistin (COL) responses. Black = 2.0, 1.0, and 0.5 μg/ml COL, respectively. **(A-xii),** Rifampin (RIF) responses. Black = 2.0, 1.0, and 0.5 mg/ml RIF. **B,** Representative synograms showing the impact on PA14 bacterial levels (a.u., defined as the AUCs of OD600 for 30 hours) of the presence (black) or absence (red) of sub-MIC antibiotics with (grey zone) or without (white zone) DMS3vir phage at MOI = 10. Data are shown for (**B-i**) synergistic, (**B-ii**) antagonistic, and (**B-iii**) non-additive (or independent) phage-antibiotic interactions. **C,** The KIM-C1 phage cocktail plus aztreonam mediated suppression at different thresholds of efficacy (>50, >60, and >70% suppression).

**Supplementary Table 1** **| *Pa* strain PA14 Receptor Mutations Conferring Phage-Resistance.**

| **Phage Treatment** | **Gene(s)** | **Role of Gene(s)** | **Mutation Type** | **Amino acid Change** |
| --- | --- | --- | --- | --- |
| **DMS3vir** | *pilT* | Twitching motility | Frame Shift | - |
| **TIVP-H6** | *pilN* | Twitching motility | Substitution | 61: F 🡪 I |
|  |  |  | Substitution | 58: Q 🡪 V |
|  |  |  | Substitution | 56: Q 🡪 L |
|  |  |  | Substitution | 50: RK 🡪 QF |
|  |  |  | Substitution | 47: Q 🡪 L |
|  |  |  | Substitution | 43: E 🡪 K |
| **Luz24** | *pilD* | Twitching motility | Substitution | 2: E 🡪 P |
|  | *pilF* |  | Substitution | 243: P 🡪 L |
|  | *pilK* |  | Substitution | 315: V 🡪 A |
|  | *pilJ* |  | Substitution | 140: A 🡪 V |
| **PAML-31-1** | *wapR* | Rhamnosyltransferases involved in LPS CPA and OSA biosynthesis | Substitution | 85: A🡪 T |
|  | *wbpZ* | Rhamnosyltransferases involved in LPS CPA and OSA biosynthesis | Substitution | 338: A 🡪 T |
|  | *migA* | LPS core biosynthesis | Substitution | 89: M 🡪 V |
| **LMA2** | *wapR* | Rhamnosyltransferases involved in LPS CPA and OSA biosynthesis | Substitution | 78: K 🡪 R |
|  | *wbpZ* | Rhamnosyltransferases involved in LPS CPA and OSA biosynthesis | Substitution | 181: A 🡪 T |
|  | *migA* | LPS core biosynthesis | Substitution | 275: Q 🡪 R |
|  |  |  | Substitution | 508: M 🡪 V |
| **OMKO1** | *flgL* | Swimming motility | Frame Shift | - |
|  | *oprM* | Multidrug efflux and transporter protein | Substitution | 50: V 🡪 G |

**Supplementary Table 2** **| Detailed information on the phage cocktails used shown in Fig 4A.** Suppression Index measured for 1st Phage (cocktails). Resistance Index from PA14 that was isolated after 30 hours-exposure from the1st Phage (cocktails) treatment, and subsequently re-challenged with each constituent of the cocktail pattern as indicated, each with MOI = 100.

| **Groups** | **1st Phage Treatment**  **(for Suppression Index shown in Fig 4A-ii)** | **2nd Phage Treatment**  **(for Resistance Index shown in Fig 4A-iii)** |
| --- | --- | --- |
| **1**Φ | DMS3vir | DMS3vir |
|  | TIVP-H6 | TIVP-H6 |
|  | Luz24 | Luz24 |
|  | 14-1 | 14-1 |
|  | phiKZ | phiKZ |
|  | PAML-31-1 | PAML-31-1 |
|  | LMA2 | LMA2 |
|  | OMKO1 | OMKO1 |
| **2**Φ  **Same CG** | DMS3vir + TIVP-H6 | DMS3vir |
|  |  | TIVP-H6 |
|  | DMS3vir + Luz24 | DMS3vir |
|  |  | Luz24 |
|  | TIVP-H6 + Luz24 | TIVP-H6 |
|  |  | Luz24 |
|  | 14-1 + phiKZ | 14-1 |
|  |  | phiKZ |
|  | PAML-31-1 + LMA2 | PAML-31-1 |
|  |  | LMA2 |
| **2**Φ  **Diff. CGs** | DMS3vir + 14-1 | DMS3vir |
|  |  | 14-1 |
|  | DMS3vir + LMA2 | DMS3vir |
|  |  | LMA2 |
|  | DMS3vir + OMKO1 | DMS3vir |
|  |  | OMKO1 |
|  | 14-1 + PAML-31-1 | 14-1 |
|  |  | LMA2 |
|  | 14-1 + OMKO1 | 14-1 |
|  |  | OMKO1 |
|  | LMA2 + OMKO1 | LMA2 |
|  |  | OMKO1 |
| **3**Φ  **Diff. CGs** | DMS3vir + 14-1 + LMA2 | DMS3vir |
|  |  | 14-1 |
|  |  | LMA2 |
|  | DMS3vir + 14-1 + OMKO1 | DMS3vir |
|  |  | 14-1 |
|  |  | OMKO1 |
|  | DMS3vir + LMA2 + OMKO1 | DMS3vir |
|  |  | LMA2 |
|  |  | OMKO1 |
|  | 14-1 + PAML-31-1 + OMKO1 | 14-1 |
|  |  | PAML-31-1 |
|  |  | OMKO1 |
|  | Luz24 + PAML-31-1 + OMKO1  (= **Cocktail KIM-C1**) | Luz24 |
|  |  | PAML-31-1 |
|  |  | OMKO1 |

**Supplementary Table 3 | Bacterial strains**

| **Lab strains** | **Genotype/description** | **References/ Sources** |
| --- | --- | --- |
| ***P. aeruginosa*** | | |
| PA14 | UCBPP-PA14m, a human clinical isolate | [1]/ Gitai Lab |
| PA14-*gfp* | PA14 *attTn7::[PA1/04/03-gfp]* | [1]/ Gitai Lab |
| PA14-*mCherry* | PA14 *attTn7::[PA1/04/03-mCherry]* | [1]/ Gitai Lab |
| PA14*ΔpilA* | PA14 *ΔpilA aacc1::FRT* | [2]/ Gitai Lab |
| PA14*ΔpilB* | PA14 *ΔpilB aacc1::FRT* | [2]/ Gitai Lab |
| PA14*ΔpilC* | PA14 *pilC::Tn5* | [3]/ Gitai Lab |
| PA14*ΔpilE* | PA14 Unmarked in-frame deletions of *pilE* | [4]/ Gitai Lab |
| PA14*ΔpilW* | PA14 Unmarked in-frame deletions of *pilW* | [4]/ Gitai Lab |
| PA14*ΔpilTU* | PA14 *ΔpilTU::FRT* | [1]/ Gitai Lab |
| PA14*ΔpilX* | PA14 Unmarked in-frame deletions of *pilX* | [4]/ Gitai Lab |
| PA14*ΔpilY1* | PA14 *ΔpilY1 aacc1::FRT* | [2]/ Gitai Lab |
| PA14*ΔfliC* | PA14 *ΔfliC aacc1::FRT* | [2]/ Gitai Lab |
| PA14*ΔflaK* | PA14 *flgK::Tn5* | [1]/ Gitai Lab |
| PA14*ΔoprM* | PA14 *oprM::MAR2xT7* | [5]/ Hancock Lab |
| PA14*ΔwapR* | PA14 *wapR::MAR2xT7* | [6]/ Hancock Lab |
| PA14*Δ*CRISPR | PA14 *Δ*CRISPR region | [7]/ O'Toole Lab |
| PA14*ΔlasRΔrhlR* | PA14 *ΔlasRΔrhlR* | [8]/ Bassler Lab |
| PAO1 | mPAO1, a wild type | [9]/ Secor Lab |
| PAO1*∆pf4∆pf6* | mPAO1 *∆pf4∆pf6* | [9]/ Secor Lab |
| PAO1-*gfp* | mPAO1, pUCP-gfp | [10]/ Hancock Lab |
| *P. aeruginosa* Br257 | *P. aeruginosa* Br257, an environmental isolate | [11]/ Lavigne Lab |
| *P. aeruginosa* Li010 | *P. aeruginosa* Li010, a hospital sewage sample | [11]/ Lavigne Lab |
| *P. aeruginosa* GHB15 | *P. aeruginosa* GHB15, an environmental isolate | [11]/ Lavigne Lab |
| CPA001 to CPA153 | 153 human clinical isolates from Stanford Health Care | This study |
| ***S. aureus*** | | |
| *S. aureus* 1203 | *S. aureus* FB1203 | Felix Biotechnology, Inc. |
| CSA001 to CSA015 | 15 human clinical isolates from Stanford Health Care | This study |

**Supplementary Table 4 | Bacteriophage strains**

| **Phage strains (a host)** | **Lineage/description** | **References/ Sources** |
| --- | --- | --- |
| ***P. aeruginosa* Lytic Phages** | | |
| DMS3vir  (*Pa* PA14) | Duplodnaviria; Heunggongvirae; Uroviricota; Caudoviricetes; Casadabanvirus;  A lytic mutant of *P. aeruginosa* Phage DMS3 | [11]/ O'Toole Lab |
| Luz7  (*Pa* Br257) | Duplodnaviria; Heunggongvirae; Uroviricota; Caudoviricetes; Schitoviridae; Migulavirinae; Luzseptimavirus; | [12]/ Lavigne Lab |
| Luz19  (*Pa* PAO1 *∆pf4∆pf6*) | Duplodnaviria; Heunggongvirae; Uroviricota; Caudoviricetes; Autographiviridae; Krylovirinae; Phikmvvirus; | [12]/ Lavigne Lab |
| Luz24  (*Pa* Li010) | Duplodnaviria; Heunggongvirae; Uroviricota; Caudoviricetes; Bruynoghevirus | [12]/ Lavigne Lab |
| LMA2  (*Pa* GHB15) | Duplodnaviria; Heunggongvirae; Uroviricota; Caudoviricetes; Pbunavirus; Pbunavirus LMA2 | [12]/ Lavigne Lab |
| PAML-31-1  (*Pa* PAO1 *∆pf4∆pf6*) | Duplodnaviria; Heunggongvirae; Uroviricota; Caudoviricetes; Pbunavirus; Pbunavirus SN  >99% identical to *P. aeruginosa* Phage NP3 | Felix Biotechnology, Inc. |
| Terra  (*Pa* PAO1 *∆pf4∆pf6*) | Unidentified. | Felix Biotechnology, Inc. |
| TIVP-H6  (*Pa* PA14) | Unidentified. | Felix Biotechnology, Inc. |
| 14-1  (*Pa* Li010) | Duplodnaviria; Heunggongvirae; Uroviricota; Caudoviricetes; Pbunavirus; Pbunavirus pv141 | [12]/ Lavigne Lab |
| phiKZ  (*Pa* PAO1 *∆pf4∆pf6*) | Duplodnaviria; Heunggongvirae; Uroviricota; Caudoviricetes; Phikzvirus; Phikzvirus phiKZ | [12]/ Lavigne Lab |
| OMKO1  (*Pa* PAO1 *∆pf4∆pf6*) | Duplodnaviria; Heunggongvirae; Uroviricota; Caudoviricetes; Phikzvirus; unclassified Phikzvirus | Felix Biotechnology, Inc. |
| KOR_P1  (*Pa* PAO1 *∆pf4∆pf6*) | Unidentified. | This study |
| KOR_P2  (*Pa* PAO1 *∆pf4∆pf6*) | Unidentified. | This study |
| ***S. aureus* Lytic Phages** | | |
| *S. aureus* bacteriophage K  (*Sa* 1203) | Myoviridae | ATCC (ATCC 19685-B1) |
| vFB009  (*Sa* 1203) | Unidentified. | Felix Biotechnology, Inc. |
| vFB433  (*Sa* 1203) | Unidentified. | Felix Biotechnology, Inc. |
| vFB468  (*Sa* 1203) | Unidentified. | Felix Biotechnology, Inc. |
| Romulus  (*Sa*1203) | Twortlikevirus | [12]/ Lavigne Lab |
